# Two routes to land: Genomic underpinnings of parallel aerial egg deposition in aquatic Old-World *Pila* and New-World *Pomacea* (Ampullariidae)

**DOI:** 10.1101/2025.11.03.686255

**Authors:** Yufei Zhou, Huawei Mu, Xueying Nie, Yue Gao, Hui Wang, Ling Fang, Tiangang Luan, Monthon Ganmanee, Jian-Wen Qiu, Jin Sun, Jack Chi-Ho Ip

## Abstract

The parallel evolution of aerial oviposition in Old-World *Pila* and New-World *Pomacea* apple snails—diverged since the Gondwanan breakup yet convergently transitioning from aquatic to terrestrial egg deposition—offers a powerful model for probing genomic adaptations underpinning key evolutionary innovations. We generated chromosomal-level genomes of *Pila celebensis* and *Pila pesmei*, revealing a genus-specific doubling in genome size driven by transposable element expansions. Parallel chromosomal rearrangements linked to environmental sensing, stress responses, and metabolism were identified in both lineages. Gene family analyses uncovered parallel expansions in cellulases, immune genes, and environmental sensors, alongside convergent positive selection in aquaporins critical for aerial osmoregulation. Strikingly, proteomics of egg perivitelline fluid (PVF) highlighted parallel adaptations in the PV1 protein family, where increased hydrophobicity enhances desiccation resistance in aerially deposited eggs. Phylogenetic and structural evidence traced PV1’s likely origin to horizontal gene transfer (HGT) from a virus in the Ampullariidae ancestor, with subsequent duplications enabling lineage-specific aerial adaptations. Despite distinct genomic trajectories, *Pila* and *Pomacea* demonstrate that parallel molecular evolution and HGT-driven innovation facilitated their convergent transition to terrestrial reproduction, providing unprecedented insights into the genomic architecture of adaptive convergence.

## Introduction

Evolutionary transition from water to land, termed terrestrialization, represents one of the most remarkable events that have profoundly shaped Earth’s biodiversity (Romans-Palacios et al., 2022). In animals, this shift occurred independently across at least nine phyla (chordates, nematodes, tardigrades, arthropods, onychophorans, platyhelminthes, molluscs, annelids, and nemerteans) over the past 420 million years (Martínez-Redondo et al., 2023). A core biological question is how genomic adaptations responded to the distinct physical and ecological challenges during the transition to land. In vertebrates, several mechanisms have been proposed, including: (i) transposable element (TE) proliferation, which rewired genome structure and regulation, enabling innovations like limb development (Falcon et al., 2023); (ii) gene family dynamics and co-option, generating novel genetic material for traits such as aerial respiration and desiccation tolerance (Meyer et al., 2021; Wang et al., 2021a; and (iii) novel regulatory networks, such as Hox cluster linkages for the fin-to-limb transition, as observed in lungfish (Meyer et al., 2021). In invertebrates, genomic studies have also provided insights into terrestrialization. For example, Vargas-Chávez et al. analyzed chromatin structures in marine, freshwater, and terrestrial annelids, while Martínez-Redondo et al. investigated ∼100 mollusc genomes to explore independent transitions from marine to terrestrial environments (Vargas-Chávez et al., 2024; Martínez-Redondo et al., 2023). These studies offer a foundation for understanding the genomic basis of terrestrial adaptations in diverse taxa.

Gastropods (Mollusca) are exceptional models for dissecting invertebrate terrestrialization genomics, having independently colonised land at least eight times, often via freshwater or brackish routes (Krug et al., 2022; Vermeij et al., 2022). Many terrestrial snails have evolved behavioral and metabolic adaptations to maintain thermoregulation and water balance, such as seeking shaded habitats, burrowing into soil, and modulating shell size and pigmentation (Staikou, 1999; Yom-Tov, 1971), whereas some species also have specialized mantle opening for aerial respiration (Krug et al., 2022; Knigge et al., 2017). At the cellular level, terrestrial snails exhibit an enrichment in chaperones and antioxidant enzymes, alongside responses in mucocyte and digestive gland cells, facilitating their transition to land (Schweizer et al, 2019). Within gastropods, the apple snails (Ampullariidae) serve as ideal models for understanding the transition from water to land (Sun et al., 2019; Haye et al., 2009), similar to the role of lungfish in tetrapod evolution. This family is phylogenetically significant, being basal to caenogastropods and a sister clade to land snails (Cyclophoroidea) and freshwater snails (Viviparidae) within Architaenioglossa (Barker, 2001), with a crucial trait for the terrestrialization – their ability to retain an aquatic adult life while evolving aerial oviposition (Sun et al., 2019; Haye et al., 2009). Like the evolutionary transition of insects from aquatic to terrestrial environments, changes in egg structure represent a key adaptation (Vargas et al., 2021).

Currently, Ampullariidae comprise nine genera with approximately 120 valid species distributed in almost all freshwater ecosystems in tropical and subtropical regions of Africa, America, and Asia (Cowie, 2015; Hayes et al., 2015). Within this family, the most diverse genera Old-World *Pila* and New-World *Pomacea* have undergone a remarkable parallel evolutionary trajectory, with both independently transitioning from underwater to aerial egg-laying, unlike their underwater-ovipositing sister genera (Fig. 1B) (Annate, 2024; Hayes et al., 2015; Hayes et al., 2009). Although the aerial-ovipositing ways in *Pila* and *Pomacea* are still very diverse. For example, *Pila celebensis* lays eggs in moist mud, *Pila pesmei* lays eggs on hard branches, and *Pomacea* species all lay eggs on hard substrates above the water (Fig. 1B). Adaptation to aerial oviposition necessitates overcoming core terrestrial challenges, including desiccation, osmoregulation, UV exposure, and novel predation/pathogen pressures (Selden, 2001), which are reflected in evolved traits like calcareous eggshells and specialized perivitelline fluid proteins (PVFs; Hayes et al., 2009; Heras et al., 2008). Thus, *Pila* and *Pomacea* offer a unique comparative framework: two phylogenetically distinct lineages (split after the breakup of Gondwana) parallelly evolving a critical terrestrialization trait (aerial reproduction) while otherwise remaining aquatic. Here, we leverage comparative genomics of these genera to elucidate the genomic architecture underpinning their parallel phenotypic terrestrial reproduction.

**Fig 1.**
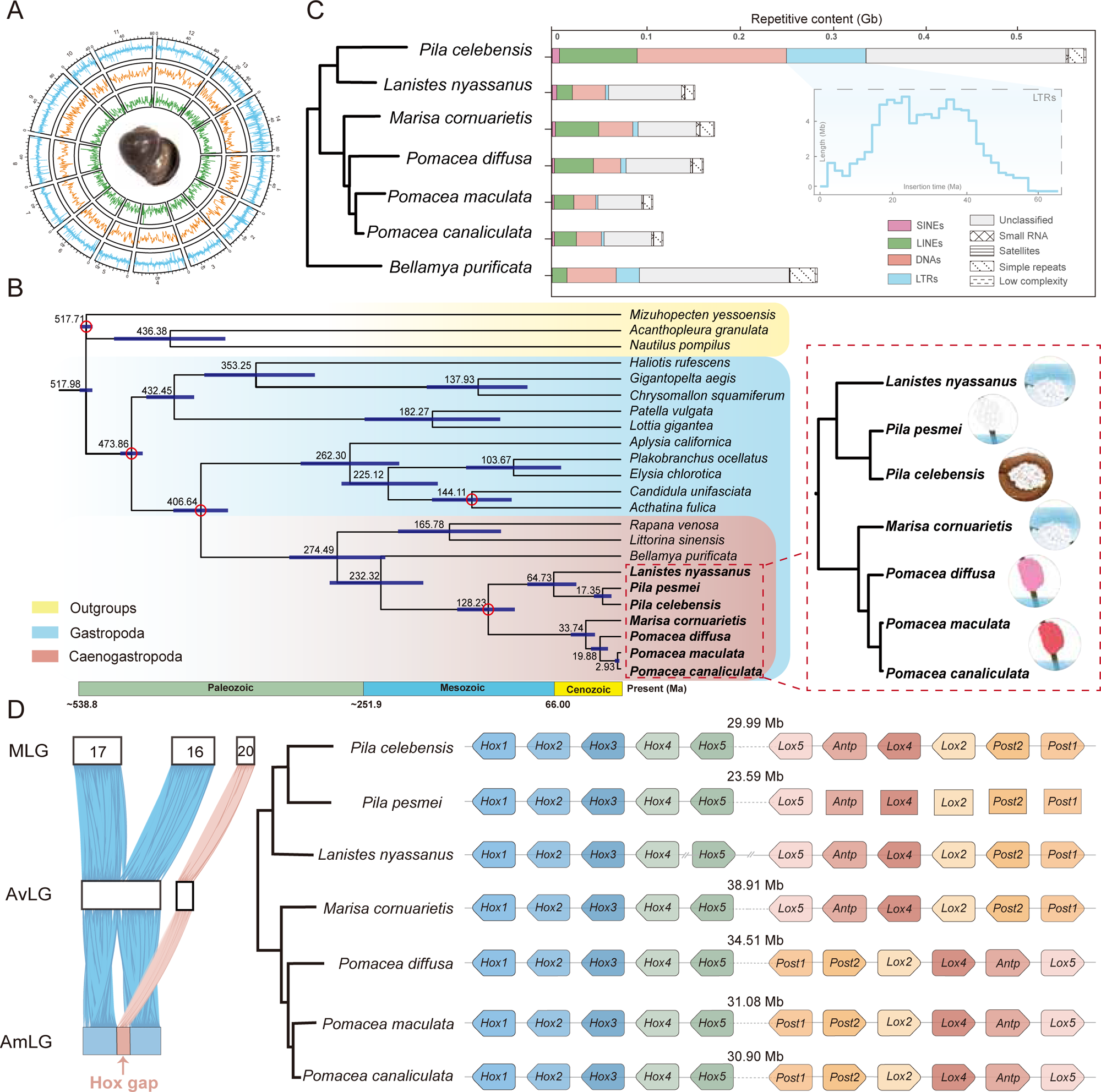
Genome landscape and evolutionary dynamics in *Pila*. **(A)** Circos plot of 14 linkage groups (corresponding to “chromosome”) showing marker distributions in 1 Mb sliding windows from outer to inner circle: Illumina sequencing depth, GC content and gene density, with the shell shape of *Pila celebensis* at the center. **(B)** Maximum-likelihood phylogenetic tree of 23 molluscs, including cephalopoda, Bivalvia, and polyplacophora as outgroups (left). The tree was calibrated at five nodes (indicated by red dots) using fossils and geological events, revealing divergence times with 100% bootstrap support; dark-purple lines indicate 95% confidence intervals. The light blue and orange branches in the Ampullariidae tree (right) illustrate aerial egg laying in *Pila* and *Pomacea*. **(C)** Comparison of repetitive elements showing expansion of four transposable element classes (i.e., SINEs, LINEs, LTRs, and DNAs) in the Ampullariidae and *Bellamya purificata* genomes. Estimated LTR insertion times for *Pila celebensis* are indicated within a gray dotted border. **(D)** Schematic of *Hox* gene clusters in seven ampullariid species, illustrating *Ho*x-linked chromosomes evolution. Gene names are labeled centrally, with connections indicating scaffolds; line lengths do not represent sequence lengths. The symbol “//” indicates a break between different scaffolds, and transcription direction is indicated by a bullet head. Ancestral linkage groups are denoted as MLG (Mollusca), AvLG (Ampullariidae and Viviparidae), and AmLG (Ampullariidae).

Our previous genomic studies have generated genome resources for studying ampullariids (i.e., genera of *Lanistes*, *Pomacea*, and *Marisa*) (Xiong et al., 2025; Ip et al., 2019; Sun et al., 2019). While these studies highlighted the role of egg neurotoxin and calcareous-related protein in New-World Canaliculata clade (*Pomacea canaliculata* and *Pomacea maculata*), the genetic basis of aerial reproduction remains largely unexplored in Old-World *Pila* and in the context of parallel evolution between Old and New World ampullariids. To fill this knowledge gap, we generated a chromosomal-level genome assembly of *Pila celebensis* and a scaffold-level assembly of *Pila pesmei*, along with a characterisation of *Pila* egg perivitelline fluid (PVF) proteome. Our analysis of parallel-evolving aerial reproductive genera reveals genome-wide gene evolution dynamics, providing foundational insights into mechanisms that have enabled the water-to-land transition in molluscs.

## Results and discussion

### Assembly and Analysis of Two Pila Genomes Reveal Repetitive Sequence Expansion Driving Genome Doubling

K-mer analysis of Illumina reads (Table S1) estimated the genome sizes of 921.5 Mb for *Pila celebensis* and 925.8 Mb for *Pila pesmei*, approximately twice the size of Old-World *Lanistes* and New-World *Marisa* and *Pomacea* spp. (Xiong et al., 2025; Sun et al., 2019). Assembly of PacBio HiFi reads resulted in 406 scaffolds for *Pila celebensis,* with an N50 of 9,853 Kb and a mean length of 2,578 Kb. Incorporating Hi-C data, 96.60% of the total genome assembly was anchored to 14 chromosomes, with the final genome size of 1,048 Mb (Figure 1A; Table S2). BUSCO analysis against metazoan_db10 BUSCOs showed 95.80% complete of *Pila celebensis*, which is comparable with newly published ampullariid and other molluscan genomes. Due to the limited samples, a hybrid assembly of Illumina reads and ONT long reads produced a draft *Pila pesmei* genome with a total length of 896 Mb (Table S2). Although the *Pila pesmei* genome has a relatively low assembly score of 87.70% metazoan BUSCOs, the high mapping rate of >98% for Illumina short reads and >90% ONT reads indicate the integrity of the assembly (Table S1). In addition, a total of 24,954 and 29,777 protein-coding genes (PCGs) were predicted in the genome of *Pila celebensis* and *Pila pesmei* with BUSCO scores of 97.20% and 86.70%, respectively (Table S2).

Synonymous substitution rate (Ks) analysis revealed no whole-genome duplication in *Pila celebensis* (Fig. S1) (Roelofs et al., 2020). To explain the doubled genome size of *Pila*, we compared repeat content across six ampullariids and one Viviparidae (Fig. 1C). *Pila celebensis* exhibits substantially higher repetitive content (577.3 Mb, 55.08% of 1.05 Gb genome) than *Lanistes* (153.7 Mb, 30.14% of 510 Mb genome), *Marisa* (174.0Mb, 32.24% of 534 Mb genome), and *Pomacea* species (∼148.0 Mb, ∼27.06% of ∼459 Mb genomes; Table S3).

Critically, non-repetitive DNA is conserved across taxa (302.9–472.7 Mb), confirming repetitive element expansion drives genome size increase in *Pila*. Analysis of transposable elements (TEs) showed distinct expansion dynamics in *Pila* (Fig. 1C; Fig. S2). Long terminal repeats (LTRs: 85.0 Mb vs. 3.7 Mb average) and long interspersed nuclear elements (LINEs: 73.2 Mb vs. 27.5 Mb) underwent exceptional expansion. Such expansions haven been shown functionally significant in animals and plants: in the limestone-dwelling land snail *Oreohelix idahoensis*, LTRs, which comprise over half of its genome, contribute to genomic features through expansion, gene composition changes, and ectopic recombination (Linscott et al., 2022). In the Spanish slug *Arion vulgaris*, LINEs account for one-third of its genome, contributing to its invasive plasticity and stress resistance (Chen et al., 2020). Similarly, LTRs bursts in cotton *Gossypium rotundifolium* not only increase genome size and intergenic spacing but also affects higher-order chromosomal structures (Wang et al., 2021). Thus, the expansion of LINEs and LTRs in *Pila* may enhance environmental adaptability by remodelling gene structures or generating alleles through ectopic recombination.

### Gondwanan Breakup–Driven Global Divergence and Adaptive Evolution of Ampullariidae Revealed by Macrosynteny Analysis

We constructed a maximum-likelihood phylogenetic tree of seven ampullariids and 16 other molluscs using 7,716 orthologous genes(Figure 1B; Table S2). The topology, with full bootstrap support (100) for all nodes, recovered major Ampullariidae lineages— Caenogastropoda contains Ampullariidae and as sister group to Heterobranchia— consistent with our prior phylogenetic hypotheses (Sun et al., 2019). Fossil-calibrated divergence estimates the Old-World ampullariid lineage (*Lanistes* and *Pila*) and New-World ampullariid lineage (*Marisa* and *Pomacea*) split at 128.2 Ma (95% confidence interval of 101.5 – 155.5 Ma), aligning with Gondwana breakup at ∼120 Ma (Jokat et al., 2003). Within the Old-World lineage, *Lanistes* and *Pila* diverged ∼64.7 Ma (45.4 – 91.0 Ma), likely facilitated by land-bridge dispersals during Indian-African plate convergence (Late Cretaceous to Palaeocene; 60-70 Ma; Sil et al., 2020). The New-World lineage split between *Marisa* and *Pomacea* occurred ∼33.7 Ma (24.6 – 46.2 Ma), potentially initiated by Eocene Andean orogeny (40–50 Ma). This timing coincides with Miocene ecological shifts in Amazonia’s Lake Pebas, a proposed cradle for freshwater mollusc evolution (Hoorn et al., 2010; Wesselingh et al., 2001). In *Pomacea*, the Bridgesii clade (*P. diffusa*) and Canaliculata clade (*P. canaliculata* and *P. maculata*) diverged ∼19.9 Ma (13.5 – 28.2 Ma), while *P. canaliculata* and *P. maculata* split ∼3.0 Ma (1.9 – 4.7 Ma) – consistent with the documented introgressive hybridization between two species (Matsukura et al., 2013).

Macrosynteny analysis provides insights into species adaptation (Lewin et al., 2024). The genome of *Pila* exhibits a conserved haploid karyotype (n=14), similar with other ampullariid species, but also reveals five key fusion events from MLGs, consistent with New World species (Fig. S3 and S4). Genes involved in these chromosomal rearrangements include those related to environmental sensing and environmental adaptation (Table S4), such as regulators of G-protein signaling, where G-protein-coupled receptors (GPCRs) mediate chemical signal perception in aquatic environments, and rearrangements of synaptotagmin genes, known to enhance neural circuit complexity and learning behaviors in *Aplysia* (Orvis et al., 2022). Additional rearranged genes are associated with signal regulation, energy metabolism, and physiological activities (Table S4), further supporting the hypothesis that chromosomal rearrangements have facilitated the evolutionary adaptation of Ampullariidae to complex habitats.

In addition to environmental adaptation, we identified a fusion event that significantly altered the arrangement of *Hox* genes in Ampullariidae. All seven ampullariidae genomes retain the ancestral molluscan complement of 11 *Hox* and three *ParaHox* genes, only *Pomacea* lineage shows an inversion from *Lox*5-*Post*1 to *Post*1-*Lox*5 (Fig. S5). Specifically, a complex fusion involving MLG17, MLG16, and MLG20 (Fig. 1D) led to a ∼30 Mb inter-cluster gap between two *Hox* clusters in Ampullariidae. MLG16 and MLG17 underwent a fusion event from MLG to the common ancestor of Ampullariidae and Viviparidae, while MLG20 remained independent.

Subsequently, in the ancestor of Ampullariidae, MLG20 inserted into the middle of fused MLG16/MLG17 region, generating the ∼30 Mb gap between the two *Hox* clusters (Fig. 1D). This gap region harbors numerous genes with functional roles in energy metabolism (e.g., glyceraldehyde-3-phosphate dehydrogenase and succinate dehydrogenase), gene expression regulation, protein synthesis, and signal transport, many of which show high expression levels (Table S4), suggesting that the fusion event may have remodeled regulatory landscapes to enhance environmental responsiveness. In *Corbicula* clams, salinity stress caused significant downregulation of genes on fused chromosomes, suggesting that fusion events may influence gene expression under environmental challenges (Wang et al., 2017).

### Parallel Chromosomal Rearrangements in Pila and Pomacea

To investigate the impact of chromosomal rearrangements on the behavioral patterns of Ampullariidae species, we reconstructed the ancestral linkage group (AmLG) of the Ampullariidae common ancestor and the last common ancestor of New World species. Syntenic analyses were conducted between AmLG and *Pila celebensis*, as well as between New-World lineage MLGs and *P. canaliculata* or *M. cornuarietis* showed that chromosome numbers remained conserved across the entire family and its two ancestral nodes, despite the presence of significant inversions (Fig. 2A). Specifically, (i) Chr9, Chr8, and Chr6 of *Pila celebensis* exhibited inversions relative to AmLG; (ii) Chr2, Chr3, and Chr11 of *P. canaliculata* showed inversions compared to New-World lineage MLG; (iii) inversions were also observed in Chr2, Chr6, Chr7, and Chr12 of *P. canaliculate* when comparing *P. canaliculata* and *M. cornuarietis*. Notably, Chr2, Chr3, and Chr13 of ancestor node underwent consistent rearrangements in both *Pila celebensis* versus AmLG and *P. canaliculata* versus New-World lineage MLG, as well as not rearrangement between *M. cornuarietis* and New-World lineage MLG. These results indicate that parallel adaptive evolution may have driven these chromosomal changes in *Pila* and *Pomacea*.

**Fig 2.**
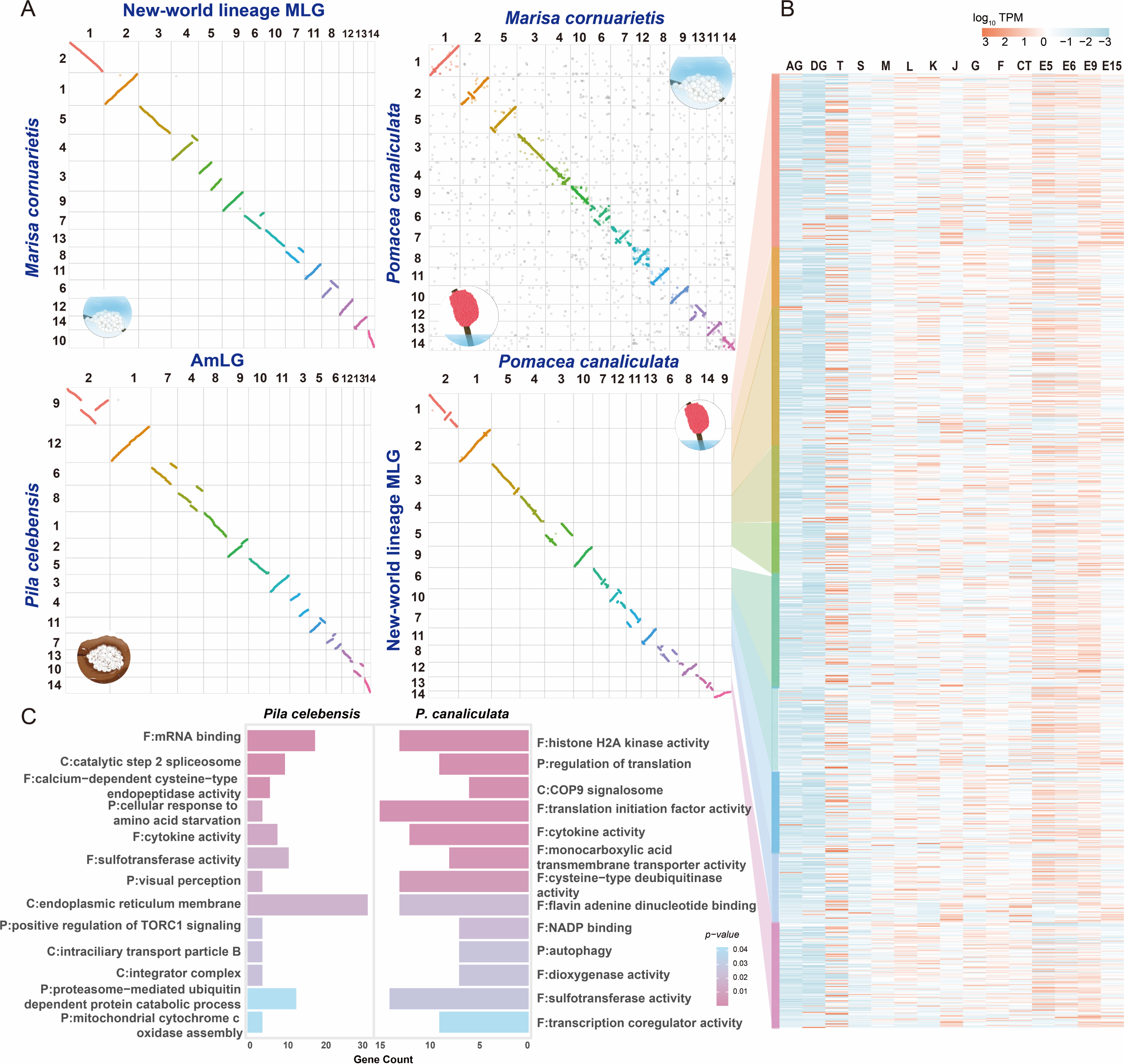
Synteny evolution and rearrangement of Ampullariidae. **(A)** Oxford dotplots of the chromosome-scale ancient gene linkage in Ampullariidae ancestor (AmLG) and New-world lineage ancestor (MLG) compared with two aerial egg-laying species (*Pila celebensis* and *P. canaliculata*) and one under water egg-laying species (*M. cornuarietis*). **(B)** Expression level of rearranged genes in different tissues in comparison of *P. canaliculata* and New-world lineage ancestor. Abbreviations of developmental stages/tissues: E5-12, day 5–15 embryo; J, juvenile; AG, albumen gland; DG, digestive gland; K, kidney; S, stomach; T, testis; L, lung; G, gill, M, mantle; F, Foot; CT, cephalic tentacle. **(C)** Gene Ontology distributions of rearranged genes in *Pila celebensis* and *P. canaliculate* compared with AmLG and New-world lineage ancestor, respectively.

Given the availability of RNAseq data from multiple developmental stages and tissues of *P. canaliculata* (Sun et al., 2019), we further examined how gene expression and associated functions relate to the rearrangements observed in the comparison of *P. canaliculata* and New-world lineage MLG (Fig. 2B). Transcriptomic analysis showed elevated expression levels during testis development and early embryogenesis (E5–E9), suggesting that chromosomal rearrangements are closely associated with early embryonic development and adaptations linked to aerial oviposition. Functional enrichment analysis revealed that genes within inversion blocks on Chr2, Chr3, and Chr13 are involved in metabolism, immune response, visual perception, and stress responses in *Pila celebensis* and *P. canaliculata* (Fig. 2C). These functions are strongly tied to the aerial oviposition process, further supporting the hypothesis that chromosomal rearrangements contributed to the parallel evolution in reproductive strategies between *Pila* and *Pomacea*. Chromosomal rearrangements can mediate environmental adaptation by altering gene function in marine mammals, functional enrichment analysis of inversion regions between two whales contains two dominant functional categories (metabolism and stress response), which are linked to cetacean adaptation, metabolism, and development (Lin et al., 2025; Zhang et al., 2020). In Gadid fishes, large-scale chromosomal rearrangements underpin speciation and cold environment adaptation (Hoff et al., 2024). Similarly, genes affected by rearrangements in both *Pila* and *Pomacea*—relative to ancestral states—are consistently associated with aerial oviposition, underscoring a convergent genomic mechanism for this reproductive innovation.

### Gene family expansion and selection shaped the genomics parallelism in aerial-ovipositing Ampullariidae

Changes in gene family size serve as a primary driver of evolution (Demuth et al., 2009). In order to capture the full breadth of genomic changes in Ampullariidae facilitating the aerial oviposition, our analysis focused on the aerial ovipositers *Pila* and *Pomacea*. To pinpoint gene repertoire changes potentially associated with the shift of in oviposition mode, we identified gene families that have rapidly expanded using BadiRate, or expanded using CAFÉ methods and positively selected in both lineages. The CAFÉ analysis revealed 197 expanded gene families in Ampullariidae, 83 expanded gene families in *Pila* and 64 expanded gene families in *Pomacea* (Table S5-S7). The BadiRate analysis detected 15 fast expanded gene families in Ampullariidae, 59 in *Pila* and 232 in the *Pomacea* (Fig. 3A; Table S8). The branch-site and site models detected a total of 1965 positively selected gene families were detected in Ampullariidae, with 1729 in *Pila celebensis* and 628 in *Pomacea* (Fig. 3A; Table S9).

**Fig 3.**
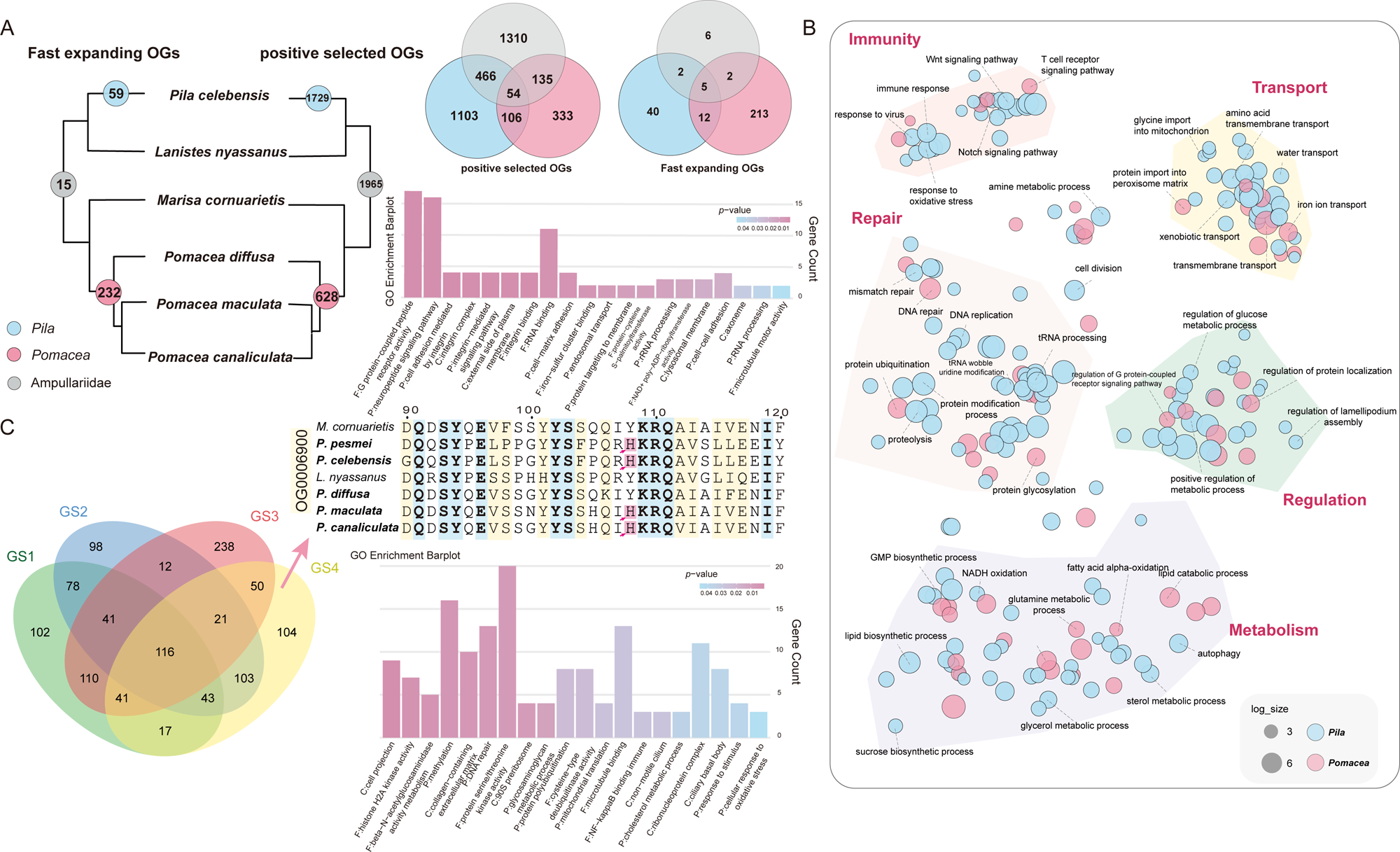
Functional characterization of parallel and exclusive orthogroup changes in *Pila* and *Pomacea* lineages. **(A)** Numbers in the colored circle indicate the count of fast-expanding and positive selected orthogroups (OGs) across different lineages. Venn diagram on the right highlights the overlapping of fast-expanding and positive selected OGs, while the Gene Ontology (GO) distributions illustrate all corresponding overlapped genes in *Pila* and *Pomace*. **(B)** Clustering based on GO enrichment results for uniquely fast-expanding and positive selected genes in *Pila* and *Pomacea*. **(C)** Convergence analysis of the physicochemical properties of amino acids, categorized into four groups. The upper right section presents results for an immune-related protein in GS4, while the lower right shows GO enrichment results for OGs exhibiting amino acid convergence.

Consistent with previous findings in Sun et al., (2019), genomic analyses of Ampullariidae reveal expansions in gene families associated with environmental sensing, metabolic adaptation, and immune regulation. As a discussion of each expanded gene family would be impossible, we here focus on discussing with more details of some representative genes in the above functions. For the metabolic adaptation, cellulase family 10 (GH10) and β-D-xylosidase exhibit significant expansion, which are key enzymes involved in the degradation of complex carbohydrates and catalyse the hydrolysis of cellulose (Table S5 and Fig. S6) (Sade et al., 2021). An increased repertoire of these kind of enzymes could have facilitated the adaptation of environmental shift, which aligned with the omnivorous diet of ampullariids including algae, plants, detritus, and zooplankton (Hayes et al., 2015). Aristide et al. also suggested that the expansion of glycosyl hydrolases could have facilitated the dietary shift associated with the habitat transition (Aristide et al., 2023). In addition, immune-related gene families also exhibit significant expansion, including C1q-related, kelch-like, and ficolin-1-like proteins. The expansion of these genes could enhance the resistance of ampullariids to environmental pathogens (Wang et al., 2012). Notably, the expansion of C1q-related proteins may be associated with the PVF evolution, which supports immune defense during embryogenesis (details in Fig. S7) (Ip et al., 2019).

Among the gene family under positive selection, we identified aquaporins (AQPs) as exhibiting parallel positive selection between *Pila* and *Pomacea* (Table S9–S12). AQPs are transmembrane transport proteins that form channels facilitating water and small solute movement across membranes and play critical roles in osmoregulation (Kruse et al., 2006). Given the stark osmotic differences between aquatic and aerial environments, the adaptive evolution of AQPs likely supports the out-of-water oviposition strategy shared by both genera (Aristide et al., 2023; Martínez-Redondo et al., 2023). Similarly, the gene families undergoing parallel positive selection in *Pila* and *Pomacea*, like the expanded gene families, are primarily involved in functions such as signal transduction, metabolism, and immunity. This parallel evolution facilitates their adaptation to the aerial oviposition process.

### Convergent Evolution of Gene Functions and Amino Acid Properties in Aerial-Ovipositing Ampullariidae

Although only one expanded gene family in the CAFÉ results and 17 fast-expanding families identified by BadiRate are shared between *Pila* and *Pomacea*, most expansions are lineage-specific and primarily associated with environmental signal perception and regulation, immune defense, and energy metabolism. (Fig. 3B; Table S6 - S7). These findings suggest that *Pila* and *Pomacea* have undergone convergent functional adaptations, utilizing similar biological pathways but different gene sets. Here, we focus on representative genes involved in environmental sensing, UV resistance, immunity, and desiccation resistance to explore their convergent adaptation mechanisms. For environmental sensing, the GPCR family, crucial for chemoreception in aquatic snails (Adema et al., 2017; Sun et al., 2019), exhibits notable expansions in *Pila* (154 genes) and *Pomacea* (31 genes), far more than in other ampullariids (3–5 genes). Phylogenetic analysis reveals *Pila* GPCRs cluster into large clades, while *Pomacea* GPCRs are more dispersed (Fig. S8). Many GPCR genes in both genera are tandemly duplicated, with *Pila* showing more extensive tandem duplications (Fig. S9).

In UV resistance, lineage-specific expansions of UV-related gene families were observed in both genera (Fig. S10). *Pila* exhibits significant expansion of keratin genes (14 copies, compared to 1–5 in *Pomacea*), potentially enhancing cuticular UV protection (Nirmal et al., 2024). In contrast, *Pomacea* shows expansion of the neprilysin gene family (9–13 copies, compared to 3 in *Pila*).

While the function of neprilysin in molluscs remains unclear, its role in UV-responsive pathways in mammals indicates a convergent mechanism (Imokawa et al., 2015). Immunity also plays a key role in aerial oviposition. *Pila* shows significant expansions in the interleukin-1 receptor (IL- 1R) and fucolectin-3-like gene families (Table S6). IL-1R mediates immune and inflammatory responses to infections (Dinarello, 1998), while fucolectin-3-like genes are defensive agents in *Xenopus tropicalis* (Mitros et al., 2019). Conversely, *Pomacea* exhibits a marked expansion of cytochrome P450 (CYP) genes, essential for xenobiotic detoxification and stress hormone biosynthesis (Denisov et al., 2005), aligning with prior reports linking CYP expansion to stress resistance and terrestrial adaptation in molluscs (Aristide et al., 2023; Liu et al., 2018).

Additionally, positively selected genes differ between the genera. In *Pila*, genes linked to bioenergetic metabolism (e.g., hexokinase and TCA cycle genes) are under positive selection, while in *Pomacea*, genes related to oxidative stress resistance, including CYP450 and flavin-containing monooxygenase (FMO) families, show signs of adaptive evolution (Tables S11-12). These adaptations likely enhance resilience to environmental stresses.

For the analysis of amino acid (AA) properties, AAs were split into four groups depending on their properties, namely (i) GS1 is based on side chain polarity and charge; (ii) GS2 reflects hydrophobicity and functional similarity; (iii) GS3 is organized by chemical reactivity and structural roles; (iiii) GS4 fine-grained scheme uses individual physicochemical traits. In total, there are 548 OGs in GS1, 512 OGS in GS2, 629 OGs in GS3 and 495 OGs in GS4 that have AA sites with similar physical and chemical properties, and there are 116 OGs with convergent evolution of amino acid sites in all four classifications (Fig. 3C; Table S13). Functional enrichment analysis showed that the genes located in these AA sites convergent OGs are primarily involved in metabolism, immune, regulation of organic compounds as well as response to oxidative stress and stimulus. This result further supports the convergent evolution of AA sites in *Pila* and *Pomacea*. The functions of these convergently evolved OGs are also closely related to the aerial-ovipositing process of *Pila* and *Pomacea*. Overall, *Pila* and *Pomacea* exhibit functional convergence and convergent evolution of amino acid properties in high-level biological processes associated with the transition to aerial oviposition, though these adaptations were achieved via distinct genomic pathways.

### Egg proteomes show the parallel selection of PV1 subunits in aerial and aquatic eggs

Egg PVF proteins have been the primary focus in the water-to-land transition of ampullariids over the last three decades (Ip et al., 2020; Ip et al., 2019; Giglio et al., 2018; Pasquevich et al., 2014; Sun et al., 2012), but there is a lack of PVF data in Old-World species. To better understand the function and evolutionary history of PVF proteins, we generated Old-World *Pila* and *Lanistes* PVF proteomes. LC-MS/MS analysis identified 57 and 133 putative proteins in *Pila pesmei* and *Lanistes nyassanus* respectively, with 47 (82.5%) and 116 (87.2%) successfully annotated against the NCBI non-redundant database (Table S14). Most of the PVF proteins are highly expressed in albumen gland while weakly expressed in the other tissues (Table S14).

Based on the annotation of PVF proteins, PVF proteins of each species are mainly divided into several functional categories such as PV1, PV2, immunity, protein degradation, and antioxidant stress (Fig. 4A). Strikingly, homologs of *Pomcea* PV1 constituted the predominant components of PVF proteomes across ampullariids with higher relative abundance in aerial eggs (*Pila pesmei*: 90.79%; *P. canaliculata*: 84.72%; *P. maculata*: 87.0%) than these in aquatic eggs (*L. nyassanus*: 53.81%; *M. cornuarietis*: 73.2%). Transcriptomic profiling further corroborated these findings: PV1-like genes showed AG-specific overexpression (TPM > 500), while undetectable/weakly expressed in other tissues (Table S14). The tissue-restricted expression pattern and PVF dominance of PV1-like proteins strongly support their essential role in embryonic survival.

**Fig 4.**
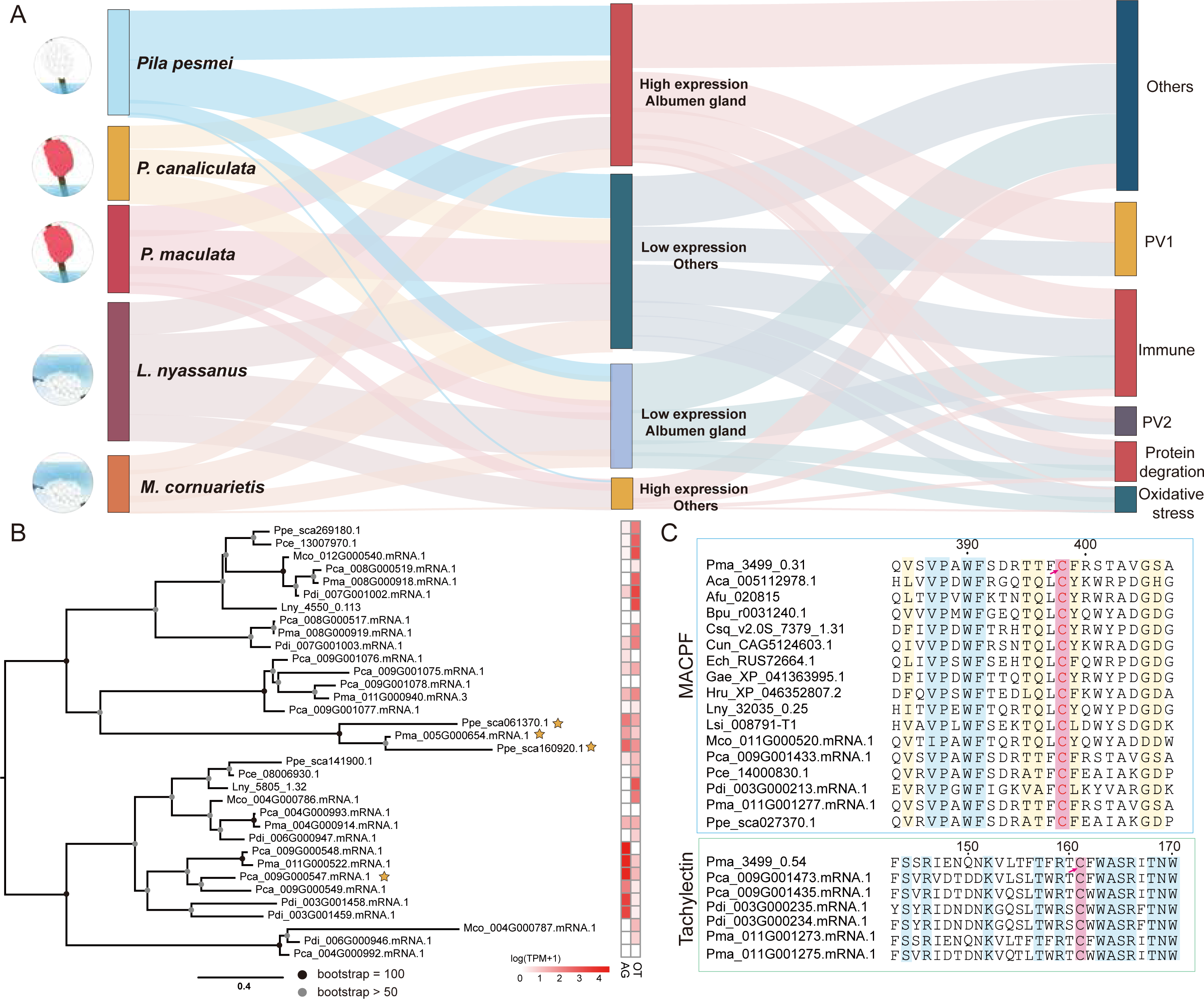
Profile of perivitelline fluid protein (PVF) in Ampullariidae. **(A)** Sankey plot of total PVF in five Ampullariidae species (*Pila celebensis*, *Pomacea canaliculata*, *Pomacea maculata*, *Lanistes nyassanus* and *Marisa cornuarietis*). The middle column shows the expression of different PVFs in the albumen gland and other tissues, and the right side shows the annotated functions of perivitelline fluid proteins. **(B)** Maximum-likelihood phylogenetic tree of calcium binding proteins (CaBP) in Ampullariidae, with circles indicating bootstrap values. Sequences marked with yellow stars were detected in the protein MS data. Gene expression levels are presented on a logarithmic scale. AG: albumen gland; OT: other tissues. **(C)** Alignment results of MACPF-like genes across all species used in Fig. 1C and tachylectin-like genes in Pomacea species. The pink background highlights the disulfide bond sites facilitating the binding and toxicity of MACPF and tachylectin.

Conversely, aquatic ovipositors exhibited enhanced immunological investment, with immune-related proteins representing 29.76% (*L. nyassanus*) and 16.8% (*M. cornuarietis*) of PVF proteomes, versus only 2.12-4.31% in aerial egg depositors (Table S14). The high abundance of immune proteins in aquatic snail eggs might reflected higher investment in immune related proteins for defence against pathogens. In addition, we observed the presence of calcium-binding proteins in aerial eggs of *Pila* and *Pomecaea*. CaBP protein was detected in the PVF protein MS of *Pila* and *Pomacea*, but there were differences in their expression levels (Fig. 4B; Table S14). Although the TPM of CaBP in *Pila* is >380, in *Pomacea*, the TPM of CaBP is >10,000. Although this result may indicate parallel evolution of CaBP in aerial ovipositioners and consistent with the hypothesis that that aquatic eggs need more protection for their embryos under the threat of microbial infection (Ip et al., 2020; Ip et al., 2019; Benkendorff et al., 2001), there are still certain differences in their evolution and expression patterns.

PV2 are novel neurotoxic PVF proteins exclusively identified in the Canaliculata clade (*P. canaliculata* and *P. maculata* (Giglio et al., 2020; Sun et al., 2019; Heras et al., 2008). This heterodimeric toxin consists of a MACPF subunit (PV2-67, 67 kDa) and a tachylectin-like subunit (PV2-32, 31 kDa) (Sun et al., 2012). Previous studies have detected a cysteine site in MACPF (Cys398) and tachylectin (Cys161). These two cysteines can interact to form a disulfide bond that connects MACPF and tachylectin together to exert toxic functions. It is believed that members of the Canaliculata lineage have produced a new toxic complex based on the function of the previously existing protein, which has a new role in defending against predators (Matías Leonel Giglio, et al., 2020). In this study, we reanalyzed the composition of PV2, with detailed in the supplementary materials and methods. We found that MACPF is conservative across most species in the species tree in Fig. 1, but tachylectin is only present in Caenogastropoda, and only in the Canaliculata lineage that tachylectin has a Cys binding site (Fig. 4C). Moreover, tachylectin is highly expressed in AG. Although *P. diffusa* also has a Cys binding site, the TPM is <1 in the albumen gland (Table S14; Table S16). This result explains why only the PVF of the Canaliculata lineage species contains the toxic protein complex PV2. This may also have contributed to the high invasiveness of some members of the Canaliculata clade (Brola et al., 2021).

PV1 is a major perivitellin that forms glyco-lipoprotein complexes in PVF, and is uniquely present in the Ampullariidae family, with no presence in the other two architaenioglossan families, Cyclophoridae and Viviparidae. BLASTp analysis identified 46 PV1-like genes across eight ampullariid species: six genes in four *Pomacea* species, seven in *Marisa cornuarietis*, six in *Lanistes nyassanus*, five in *Pila pesmei*, and four in *Pila celebensis*. Phylogenetic analysis of the PV1 genes revealed five major clades (Fig. 5A). Most clades retained the phylogenetic relationship of the species tree, except for clade V, which exhibited New World-specific gene duplications in *Pomacea* and *Marisa*. Sequence alignment and substitution analyses suggest that clades I–IV originated from ancestral duplications in ampullariids, with divergence times aligning with species divergence (Fig. S11 – Fig. S12). Clade V in New World lineages (*Marisa* and *Pomacea*) diverged around 10 million years ago, coinciding with the Andean uplift (Hoorn et al., 2010). This temporal correlation implies that environmental changes during this period may have driven rapid adaptation through gene duplication.

**Fig 5.**
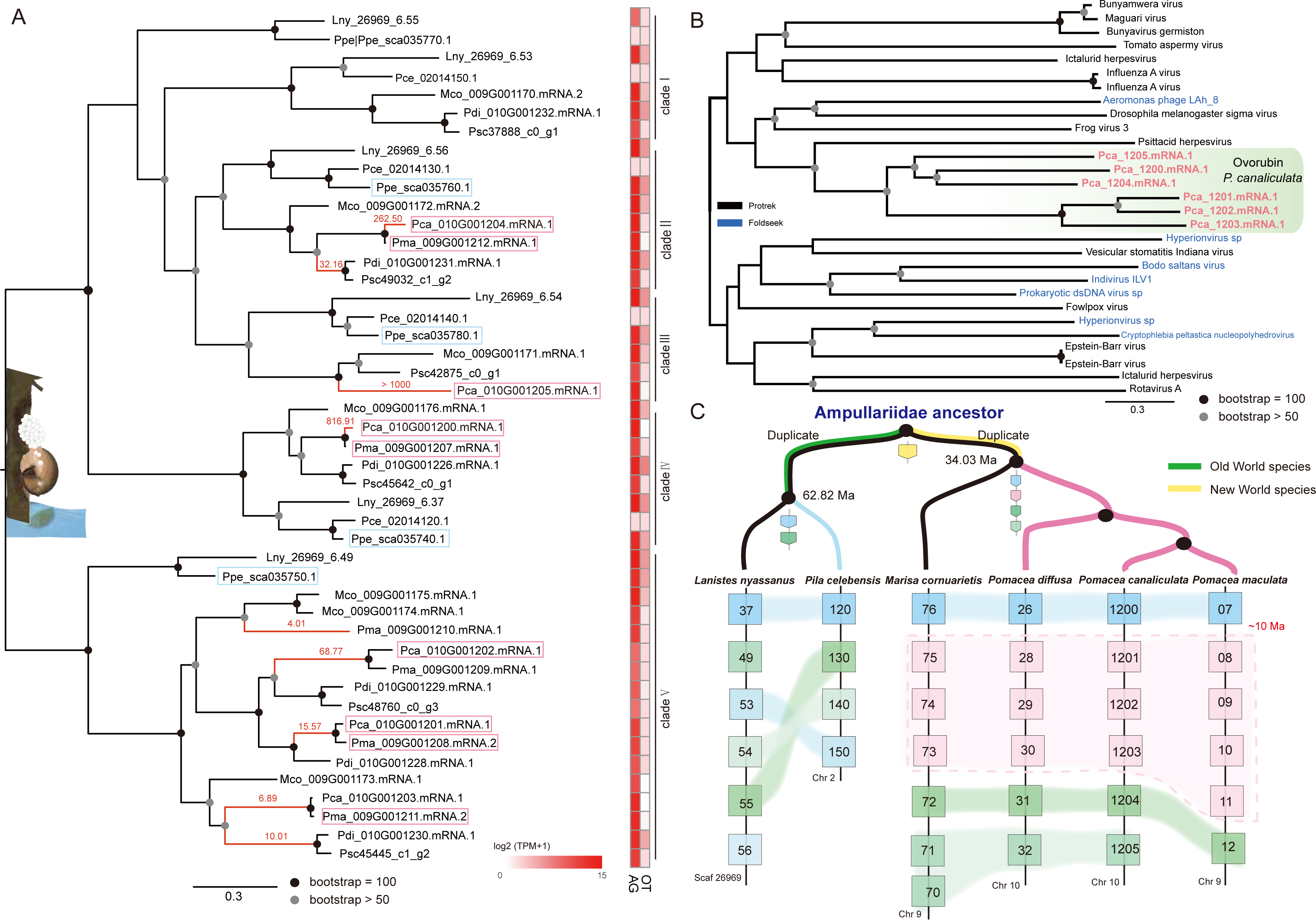
Phylogeny and evolution prediction of PV1 in Ampullariidae. **(A)** Phylogenetic tree and expression of PV1 homologues in Ampullariidae. Blue and pink frames indicate detection of proteins in the mass spectrometry (MS) dataset. Red branches signify positive selection, with dN/dS values shown in red. Gene expression levels are on a logarithmic scale. AG: albumen gland; OT: other tissues. **(B)** Phylogenetic analysis supporting the horizontal acquisition of PV1 from viruses. Sequences in pink represent six PV1 sequences from *P. canaliculata*, blue sequences are the viral sequences obtained via Foldseek through 3D structure alignment, and the others are predicted using Protrek. **(C)** Schematic representation of PV1 evolution, with the yellow section indicating the common ancestor of PV1. Each node section indicates the ancestral number of PV1, and the pink section indicates a recent copy event occurring ∼10 million years ago (Ma).

Gene duplication provides new genetic material for selection (Zhang 2003), with clade V of *Pomacea* showing evidence of positive selection (Fig. S12). Notably, positively selected sites revealed substitutions to aromatic residues, particularly phenylalanine (Phe). In aerial egg-laying species, the Phe content in clade V of PV1 is approximately double that of aquatic egg-laying species (Table S15), along with an increased hydrophobic-to-aromatic amino acid ratio. This likely enhances PV1 binding to carotenoids (Shahidi et al., 2023), improving antioxidant capacity, UV resistance, and aposematic signaling in the Canaliculata clade of *Pomacea* (Pasquevich et al., 2024; Heras et al., 2008). Furthermore, aerial ovipositors in *Pomacea* (clade V) and *Pila* (clade V) show significant enrichment in hydrophobic residues, similar with proteins in insect eggs where hydrophobic domains confer desiccation resistance (Vargas et al., 2021). The parallel evolution of hydrophobic residues in PV1 suggests a shared adaptive mechanism for protecting aerially deposited eggs from desiccation in *Pomacea* and *Pila*.

To explore the origin of PV1, we analyzed six PV1 sequences from *P. canaliculata* using Protrek and FoldSeek for 3D structural homology. Protrek identified high similarity scores (20–35) with viral sequences, while FoldSeek indicated structural similarities between PV1 and viral protein folds. Phylogenetic analysis of the top hits (p < 0.05) revealed that all PV1 sequences from *P. canaliculata* cluster closely with viral sequences, such as Psittacid herpesvirus (Fig. 5B), supporting the hypothesis that PV1 originated via horizontal gene transfer (HGT) from viruses. Advances in genomics have increasingly uncovered HGT in molluscs, with previous reports documenting viral-derived HGT in the sea slug *Elysia crispata* and scallop *Chlamys farreri* (Eastman et al., 2023; Li et al., 2025). Our study provides a notable example of viral HGT in molluscan lineages that have functional consequences.

Genomic analysis revealed at least four tandemly arranged PV1 genes on the same chromosome (Fig. 5C). Based on divergence time estimates (Fig. S11–S12) and HGT evidence, we propose a model for PV1 evolution in Old World and New World ampullariids. The ancestral PV1 gene, acquired via viral HGT (yellow branch in Fig. 5C), underwent duplication and divergence during ampullariid evolution. In Old World lineages, the ancestral PV1 gene duplicated before to the divergence of *Pila* and *Lanistes*, followed by lineage-specific duplications, resulting in six PV1 copies in *L. nyassanus* and four in *Pila celebensis*. In New World lineages, the ancestral PV1 gene formed a tandem array of four copies through duplication. Approximately 10 million years ago, after the divergence of *P. diffusa* and the Canaliculata clade (*P. canaliculata* and *P. maculata*), one copy underwent further duplication, producing seven PV1 copies in *M. cornuarietis* and six in *Pomacea* species. In summary, Ampullariidae ancestors acquired PV1 through viral HGT, with subsequent gene duplication and functional divergence enabling the evolution of both aerial and aquatic egg-laying strategies within the family.

## Conclusion

This study provides the first chromosomal-level genomes for Old-Wolrd *Pila*, offering a pivotal resource for dissecting the genomic basis of convergent evolution. We demonstrate that the independent origins of aerial oviposition in *Pila* and *Pomacea* were driven by multi-layered genomic parallelism. Macrosynteny analyses reveal deep homology, including shared ancestral chromosomal fusions that remodeled the Hox cluster and parallel inversions on homologous chromosomes (Chr2, Chr3, Chr13). The genes within these rearrangements—enriched for functions in metabolism, immunity, and stress response—show elevated expression during early embryogenesis, linking structural variation directly to adaptive phenotypes. At the gene level, we observe parallel expansions and positive selection in key functional categories like osmoregulation (e.g., aquaporins), while lineage-specific gene families converge on pathways for environmental sensing and UV resistance. This functional convergence is further supported by convergent evolution at the amino acid level across 116 orthogroups, reinforcing traits like oxidative stress and desiccation tolerance. Finally, the perivitelline fluid proteome reveals PV1, derived from an ancient viral horizontal gene transfer, as a core ampullariid adaptation. In aerial layers, PV1 underwent parallel gains in hydrophobicity and aromatic residues, enhancing desiccation resistance and carotenoid binding. This contrasts with the heightened immune investment in aquatic eggs and the lineage-specific evolution of the PV2 neurotoxin. Collectively, our findings demonstrate how convergent ecological innovations can emerge through parallel changes spanning chromosomal structure, gene repertoire, and protein composition, positioning Ampullariidae as a valuable model for studying ecological transitions.

## Methods

### Sample collection

We collected *Pila celebensis* specimens from Khanun District, Patthalung Province, Thailand, in 2019. We dissected one female *Pila celebensis* to obtain foot tissue for genome sequencing and dissected three additional females to various tissues (albumen gland, foot, mantle, and digestive gland) for transcriptome sequencing. We collected *Pila pesmei* individuals from Nong Phok District, Roi-et Province, Thailand, in 2012. From one female *Pila pesmei*, we collected foot tissue for genome sequencing and two egg masses for proteomic analysis. We also sampled two mass eggs from *Lanistes nyassanus* (snail source same as in Sun et al., 2019) as representatives of the Old-World underwater lineage for comparison with the *Pila* PVF. All samples were stored at -80°C before to downstream analysis. We confirmed the identity of both *Pila* species by DNA barcoding of the cytochrome c oxidase subunit I (COI) gene.

### Genome sequencing

Genomic DNA was extracted using the CTAB method (Porebski et al., 1997). DNA quality was evaluated, and quantity measured using agarose gel electrophoresis and a Qubit fluorometer (Thermo Fisher Scientific, MA, USA), respectively. High-quality DNA was then used for library preparation and whole-genome sequencing (details in Table S1).

For *Pila celebensis*, 1 µg of DNA was used to construct a library with a 350-bp insert size using the NEBNext DNA Library Prep Kit (New England Biolabs, MA, USA), and sequenced on an Illumina NovaSeq with paired-end 150 bp mode (PE150). Additionally, 10 µg of DNA was used to construct a HiFi SMRTbell library using the SMRTbell Express Template Prep Kit 2.0, which was sequenced on a PacBio Sequel II sequencer. High-quality HiFi reads were generated using the circular consensus sequencing (CCS) mode on the PacBio long-read platform. For Hi-C sequencing, the foot tissue was re-frozen on ice and mixed with 37% formaldehyde in serum-free Dulbecco’s modified Eagle’s medium DMEM. The tissue was then homogenised, treated with a restriction enzyme (MBOI), labelled with biotin, and repaired. DNA was extracted and purified for library preparation, aiming for a 350-bp insert size, using the NEBNext DNA Library Prep Kit. Finally, it was sequenced on an Illumina NovaSeq in PE150 mode.

For *Pila pesmei*, 5 µg of DNA was used to construct libraries with 250-bp and 500-bp insert sizes using the NEBNext DNA Library Prep Kit (New England Biolabs, MA, USA), as well as a mate-pair library with a 15 kb insert size using the Nextera Mate Pair Library Preparation Kit (Illumina, USA). These were sequenced on an Illumina HiSeq 2500 in PE150 mode. For long-read sequencing, 5 μg of genomic DNA was used to construct a long-length DNA library using the Ligation Sequencing Kit 1D (SQK-LSK109, ONT, Oxford, UK) according to the product’s instructions, and sequenced using one FLO-MIN106 R9.4 flow cell coupled to a GridION X5 sequencer (ONT, Oxford, UK). The raw reads were real-time base-called using Guppy v1.6.0 (ONT) under default settings to generate a FASTQ file.

Total RNA was extracted from three individuals of *Pila celebensis* using Trizol reagent (Thermo Fisher Scientific, MA, USA), following the manufacturer’s protocol. The quality of the RNA samples was assessed by agarose gel electrophoresis, and the quantity was measured using a Qubit fluorometer (Thermo Fisher Scientific, MA, USA). RNA samples were submitted to Novogene (Beijing, China) for cDNA library preparation using the Illumina NEBNext Ultra RNA Library Prep Kit (New England Biolabs, MA, USA) and sequenced on an Illumina NovaSeq in PE150 mode. RNA-seq data for *Pila pesmei* (named as *Pila ampullacea* in AmpuBase) were obtained from Ip et al., 2018.

### Genome assembly

Illumina short reads were trimmed to remove adaptors and low-quality reads (quality score <30, length <40 bp) using Trimmomatic v0.38 (Bolger et al., 2014). The clean Illumina reads were used to estimate the genome sizes of the two *Pila* species using GenomeScope v2.0 (Ranallo-

Benavidez et al., 2020). For *Pila celebensis*, *de novo* assembly of HiFi reads (N50 of 9.853 Mb and mean length of 2.578 Mb; Table S1) were performed using nextDenovo v2.5.2 (Hu et al., 2024) under default settings. Possible alternative heterozygous contigs were further eliminated using purge_dups v1.2.6 (Guan et al., 2020). Possible contamination sequences (e.g., microbes, fungi, algae and viruses) were removed by search using BLASTn against the NCBI database with an E-value threshold of 1e-20 and manual correction. The completeness of the final genome assembly was assessed by analyzing the Benchmarking Universal Single-Copy Orthologs (BUSCO) v5.4.5 scores against the database metazoan_odb10 under the genome mode (Simão et al., 2015). QUAST v5.2 was used to assess assembly statistics (Gurevich et al., 2013). To further scaffold the assembly at a chromosomal level, Hi-C raw reads were trimmed using Trimmomatic v0.38 to remove low-quality reads (quality score < 30, length < 40 bp). High-quality reads were then identified with HiC-Pro v2.10 (Servant et al., 2015), and duplicates were removed using the Juicer pipeline v1.5 (Durand et al., 2016). Next, genomic scaffolding was performed with the 3D *de novo* assembly pipeline for diploid genomes (Dudchenko et al., 2017). Pseudo-chromosomal linkage groups were reviewed, and corrections were made in Juicebox v1.11.08 to ensure that scaffolds within the same groups matched the Hi-C linkage characteristics (Durand et al., 2016). RaGOO v1.1 (Alonge et al., 2019) was used to scaffold the final assembly into pseudo-chromosomal linkage groups based on *Pila celebensis* genome.

For *Pila pesmei*, the FM-index Long Read Corrector (FMLRC) was used to correct the ONT raw reads using Illumina reads (Wang et al., 2018). After trimming and error correction, a total of 50 Gb Illumina and 10 Gb ONT reads (N50 of 12.3 kb and mean length of 5.7 kb; Table S1) were retained. The hybrid assembly was conducted with MaSuRCA v3.4.1 using both Illumina and ONT data (Zimin et al., 2013). Redundances were applied to eliminate alternative heterozygous contigs (Pryszcz et al., 2016). The resulting contigs were further scaffolded with ONT data using SSPACE-LongRead v1.1 (Boetzer et al., 2014). Potential contamination sequences were removed by searching with BLASTn against the NCBI database. Genome completeness and statistics were evaluated using BUSCO and QUAST, respectively.

### Genome annotation

The *Pila* genomes were annotated according to Sun et al. (2019) and Xiong et al. (2025). In brief, the genome was “soft-masked” using RepeatMasker v4.1.0 (Smit et al., 2015) with the repeat libraries of all model organisms in the Dfam version 201810, and species-specific repeat libraries in RepeatModeler2 (Flynn et al., 2020) and EDTA v2.1.0 (Ou et al., 2019). Mollusca protein sequences from Swiss-Prot and Uniprot databases and five related-species protein sequences (*P. canaliculata*, *P. maculata*, *P. diffusa*, *M. cornuarietis* and *Lanistes nyassanus*) were used in miniprot v0.14 (Li, 2023) as protein evidence. To provide transcriptomic evidence, *de novo* and genome-guided transcriptomes of two *Pila* were assembled using Trinity v2.15.1 (Haas et al., 2013) under default settings, respectively. These two transcriptomes were merged using the

PASA pipeline v2.5.3 (Haas et al., 2003) with the “--ALIGNERS blat,gmap,minimap2” setting. BRAKER3 (Gabriel et al., 2024) was used to *ab initio* predict genes in the repeat-masked genome sequences. Results from different gene predictors were integrated into a consensus weighted annotation by EVidenceModeler (EVM) v2.1.0 (Haas et al., 2008) with “PROTEIN = 10, TRANSCRIPT = 8, ABINITIO_PREDICTION = 3” and default settings for other parameters in the software. GeMoMa v1.9 (Keilwagen et al., 2018) was used for protein homology annotation, afterwards, the annotation results from EVM and GeMoMa were combined using miniprot. Then PASA was used to improve the annotated gene models by modifying gene structures and adding UTR annotations using the de novo and genome-guided transcriptome. The predicted genes were functionally annotated using Diamond BLASTp v2.1.0 (Buchfink et al., 2021) under an E-value threshold of 1e-10 against the NCBI non-redundant (nr) database. Gene functional annotation was conducted using Blast2GO Basic 6.0 (Conesa et al., 2005) and the KAAS website (https://www.genome.jp/kaas-bin/kaas_main) for Gene Ontology (GO) and Kyoto Encyclopedia of Genes and Genomes (KEGG) pathways.

#### Phylogenomics and divergence times

Phylogenetic analysis was conducted according to Liu et al. (2023) and Sun et al. (2021). Briefly, orthologous groups (OGs) among seven ampullariids and 16 molluscan genomes were inferred using OrthoFinder v.2.5.4 (Emms et al., 2019) with Diamond BLASTp v2.1.0 under “--ultra- sensitive” mode. One genome each from Bivalvia (*Mizuhopecten yessoensis*), Polyplacophora (*Acanthopleura granulata*), and Cephalopoda (*Nautilus pompilus*) was used as outgroups of Gastropoda. Only OGs with at least 80% taxon representation (i.e., at least 18 species) were used to reconstruct their phylogenetic relationships. The protein sequences were aligned using MAFFT (Katoh et al., 2002) under default settings and trimmed using BMGE v1.12, and the alignments with fewer than 20 amino acids were removed. Each alignment was used to construct “approximately maximum likelihood” tree using FastTree version 2.1.11 (Price et al., 2010). And then PhyloPyPruner v1.2.4 (https://pypi.org/project/phylopypruner/) was used to remove paralogues. Phylogenetic analysis was conducted using a maximum-likelihood method implemented in IQ-TREE v2.2.6 (Minh et al., 2020) with the “-MFP” model and partitioned alignment to compute the best-fit model of each partition and 1000 ultrabootstraps to test the topological support. In addition, divergence times were estimated using MCMCTree v4.10.3 with various time constraints: the ‘root-age’ was set as 532 MYA; a soft constraint of 520.5-530 MYA for the origin of Bivalvia (Bieler et al., 2014); a hard lower bound of 470.2 MYA and a soft upper bound of 531.5 MYA for the first appearance of Gastropoda (Benton et al., 2009); a hard minimum 390 MYA bound for the split of Caenogastropoda and Heterobranchia (Jörger, et al., 2010); a hard upper bound of 150 MYA for the split of *Lanistes nyassanus* (Sun et al., 2019); and a hard lower bound of 130 MYA for the first appearance of both the Stylommatophora and Hygrophila (Tillier, 1996).

#### Synteny analysis

Protein sequences of *Pila celebensis* and four New-World ampullariids with the gene coordinate files (.*bed* format) were used for the ancestor linkage groups and inter-species synteny analysis compared with MLGs (Sigwart et al., 2024). The ancient linkage groups of Ampullariidae (AmLGs) and *Pomacea* ancestor were identified according to methods of Schultz et al. (2023). In brief, Diamond BLASTp v2.1.0 was conducted against corresponding species under each ancestor node, obtaining reciprocal best hits among them. Then orthologous were grouped together based on whether they exist on the same set of scaffolds in each species. After removing the groups of orthologues with a less-than-significant false discovery rate (0.05), the remaining groups, which presented on the same set of chromosomes, were considered as a common ancestor of these species. Syntenic analysis between ampullariids and ancestor linkage groups were identified through reciprocal BLAST searches, and the linkage groups were further checked Fisher’s exact test using R package macrosyntR (El Hilali et al., 2023) with a significant threshold of *p*-value < 0.01. The inter-chromosomal rearrangements were visualized using Oxford-plot (https://bitbucket.org/viemet/public/src/master/CLG/). The expression levels of genes rearranged in chromosomes in different *P. canaliculata* tissues were displayed through a heat map, and the data used was PRJNA473253 published in Sun et al. (2019).

In addition, self-BLASTp was conducted to identify the paralogous genes with a threshold of E-value of 1e-5. MAFFT and KaKs_Calculator v3.0 (Zhang, 2022) were used to align sequences and calculate the number of synonymous substitutions per synonymous sites (Ks) values, respectively. The Ks distribution was plotted using ggplot package from R v4.3.1 with “density” function.

#### Hox gene analysis

Hox gene clusters were identified using BLASTp searches against previously published ampullariid genomes (Sun et al., 2019) and other molluscan genomes, with manual verification of the Hox domain against Pfam 33.1 (Mistry et al., 2021).

### Insertion times of transposable elements

To reveal the dynamics of transposable elements (TEs) in ampullariids, repetitive elements of six ampullariids were annotated with RepeatModeler2 and RepeatMasker as described in the genome annotation section (details in Table S4). TEs and repeat landscape of Kimura substitution rate (K) were extracted from RepeatMasker results using “calcDivergenceFromAlign.pl” and “createRepeateLandscape.pl” scripts, respectively. Insertion time of TEs was estimated using the equation “T = K/2r” (Kimura, 1980), where T is the insertion time, and r is the nucleotide substitution rate for ampullariids. The substitution rate of ampullariids was estimated according to Ip et al. (2021a) using a free-ratio model in the codmel script implemented in PAML v4.9 (Yang, 2007).

#### Gene family analysis

The orthofinder results were used for gene family analysis using CAFE5 v5.0.0 (Mendes et al., 2020). Significantly expanded or contracted gene families were identified with a threshold of “family-level P-value” < 0.05. For the fast-expanding gene family analysis, we reconstructed fast-expanding gene family using BadiRate v1.3.5 as in Pau Balart-García et al. (2023). GO and KEGG enrichment analyses were conducted for expanded gene families in *Pomacea* and *Pila* using clusterProfilter R package (Yu et al., 2012). To further reveal the evolution of selected gene families, maximum likelihood trees were reconstructed using IQ-TREE with settings detailed in the phylogenomic part.

#### Positive selection and convergent evolution analysis of amino acid sites

Positive selection analysis was performed on the 7,788 orthologous genes identified using phylopyprunner, and only OGs that included at least one species in *Pomacea* species or *P. celebensis* were used in the downstream analysis. Alignment of the amino acid sequences of 7,788 OGs was conducted with MAFFT, and the nucleotide sequences of these genes were aligned according to the amino acid alignment using Pal2nal. HYPHY v2.5.61 (Kosakovsky Pond et al., 2020) was used to predict positively selected sites using Mixed Effects Model of Evolution model (MEME) and the adaptive Branch-Site Random Effects Likelihood model (aBSREL). The foreground branches (*Pomacea* species and *Pila celebensis*) were assigned in aBSREL model respectively. The nonsynonymous/synonymous substitution ratio (ω = dN/dS) and *p*-value were used to detect whether a gene was subject to adaptive evolution, with ω > 1 and *p*-value < 0.05 indicating that a gene was under positive selection. To dectect convergent evolution analysis in amino acid sites, mainly referring to the method of Chen et al., amino acids are divided into four categories according to their physical and chemical properties to test whether there is convergent evolution of amino acid sites in gene families (Chen et al., 2025). GO and KEGG enrichment analyses of positively selected genes in the *Pomacea* branch and *Pila* branch were conducted using clusterProfilter R package.

#### Analysis of PV1 and PV2

The detailed methods of PVF extraction and LC-MSMS analysis are included in the SI. To further study the evolution of PV1 and PV2 genes, we conducted the BLASTp search among the nine ampullariids (*Pila celebensis, Pila pesmei, Lanistes nyassanus, M. cornuarietis, P. canaliculata, P. maculata, P. diffusa,* and *P. scalaris* (transcriptome only)) and 16 other molluscan species genomes used in the phylogenetic tree. We also sequenced and *de novo* assembled two architaenioglossans *Cyclophorus subcarinatus* (23,873 unigenes; Cyclophoridae) and *Sinotaia aeruginosa* (23,620 unigenes; Viviparidae) using Trinity for PVF analysis. The threshold for BLASTp was set with an *E*-value of 1e^-5^, pairs aligned length ≥ 30%, and aligned sequence identity ≥ 30%.

Phylogenetic trees of PV1-like, MACPF-like and tachylectin-like genes were constructed using amino acid sequences, which were aligned using L-INS-I methods with MAFFT v7.525 (Katoh et al., 2013) and trimmed with BMGE v1.12 (Criscuolo et al., 2010). IQ-TREE was used to reconstruct a phylogenetic tree using the maximum-likelihood method and the model of MFP with a bootstrap value of 1000. The RNAseq data (SRR6395538, SRR6395540, SRR6394739, SRR6394740, SRR6395701, SRR6395700, SRR6395705, SRR6395706, SRR6395702, SRR6394741, SRR6394742, SRR6394704, SRR6394703, SRR6395720 and SRR6395719) were mapped to the 6 genomes and 1 transcriptome for determination of gene expression level expressed as Transcripts Per Kilobase Million (TPM) using Salmon v1.8.0 (Patro et al., 2017).

Selective pressure was determined by the ratio (ω = dN/dS) with purifying, neutral, or positive selection indicated by ω < 1, = 1, or > 1, respectively. Substitution rate of each branch was estimated using the HYPHY to perform an exploratory analysis with the aBSREL and MEME model respectively (similar to the above section). PV1-like, MACPF-like and tachylectin-like genes were used to infer the divergence time respectively. The divergence times were estimated with BEAST v2.6.5 (Bouckaert et al., 2014). An XML (Extensible Markup Language) file was generated with BEAUti (version 2, included in the BEAST package). The three phylogenetic trees obtained from the obvious step were fixed by removing Wide-exchange, Narrow-exchange, Wilson-Balding and Subtree-slide. The rates of evolutionary changes at nuclear sites were estimated using ModelTest v3.7 with the GTR substitution model (Posada et al., 1998).

Divergence time and corresponding CIs were conducted with a log-normal relaxed molecular clock and the Yule speciation prior. Three fossil time points, i.e., a hard upper bound of 150 MYA for the split of *Lanistes nyassanus* for PV1-like genes, a hard lower bound of 470.2 MYA and a soft upper bound of 531.5 MYA for MACPF-like genes and a hard minimum 390 MYA bound for tachylectin-like genes, were selected for calibration. After 10,000,000 generations, the first 10% were removed as burn-in. The log file was checked for convergence with Tracer v1.52 (Rambaut et al., 2018). Consequently, a maximum clade credibility (MCC) tree was summarized with TreeAnnotator (version 2.6.5, included in the BEAST package), annotating clades with > 0.8 posterior probability (PP).

In addition, dN approach was used to further estimate the gene duplication time in each species. ParaAT (Zhang et al., 2012) was used to align protein sequences with gene-coding sequences. KaKs_Calculator 3.0 was used to calculate the dN/dS ratio in all aligned pairs, and the dN value between paralogous gene pairs was determined using the MYN (Modified YN) model. Divergence time was inferred using the formula “T = dN/2R”, where R is 2.67E-8 synonymous substitutions per site per year.

We searched the PV1 for possible HGT using the AI matching score and 3D structure prediction as previously described (Su et al., 2024; Van Kempen et al., 2024). After obtaining 41 PV1- associated sequences predicted by ProtTrek and FoldSeek, we aligned these sequences using MAFFT and IQ-TREE was used to reconstruct a phylogenetic tree using the maximum-likelihood method and the model of MFP with a bootstrap value of 1000.

## Data availability

The raw sequencing datasets for *Pila celebensis* and *Pila pesmei* have been deposited in the NCBI SRA database under BioProject accessions PRJNA1355022 and the mass spectrometry proteomics data have been deposited to the ProteomeXchange Consortium via the PRIDE partner repository with the dataset identifier PXD070092. Genome assemblies, annotations of two ampullariid species can be accessed via 10.6084/m9.figshare.30479291.

## Acknowledgments

This work was financially supported by the National Key Research and Development Program of China (2022YFC2601302) and the Young Taishan Scholars Program of Shandong Province (tsqn202103036). JCHI was supported by Research Grants Council (HKSAR)’s Early Career Scheme (23100224) and General Research Fund (12102623 and 13100725).

