## Supplementary Figures for "Two routes to land: Genomic underpinnings of parallel aerial egg deposition in aquatic Old-World *Pila* and New-World *Pomacea* (Ampullariidae)"

**This Supplementary Material includes:**

1. Materials and Methods

2. Supplementary Figs. S1 to S18

3. References

**1.** **Materials and Methods**

*PVF extraction and LC-MSMS analysis*

PVF analysis of two Old-World *Pila* and *Lanstis* was performed following our previous proteomic studies (Mu et al., 2017 , Ip et al., 2018). PVF was extracted from approximately 30 eggs using a fine needle, mixed with 8 M urea, and centrifuged at 12,000×g for 10 min at 4 °C to obtain the protein-containing supernatant. Protein samples were combined with a loading buffer (20% glycerol, 0.2M Tris-HCl pH 6.8, 0.05% bromophenol blue, 10mM dithiothreitol, and 10% SDS) at a 1:3 (v/v) ratio, separated by SDS-PAGE, stained with Coomassie Brilliant Blue, and destained with 1% acetic acid. Each gel was cut into eight slices based on intensity and molecular weight, further destained with 50mM NH4HCO3 in 50% methanol, washed with MilliQ water, dried, and rehydrated twice with 100% acetonitrile and 100mM NH4HCO3. In-gel digestion was performed using sequencing grade trypsin (Promega, Madison, USA) in 50mM NH4HCO3, and peptides were recovered, desalted using Sep-Pak C18 cartridges (Waters, Milford, USA), and dried in a vacuum concentrator (Eppendorf, Hamburg, Germany).

The dried fraction from each biological sample was reconstituted with 0.1% formic acid and analyzed twice using an LTQ-Orbitrap Elite coupled with an Easy-nLC (Thermo Fisher, Bremen, Germany). Peptides were separated on a C18 capillary column (Michrom BioResources, CA) over a 90-min gradient. Mass spectrometry scans were performed in the range of 350 to 1600 m/z at a resolution of 60,000 in positive mode. The five most abundant multiple-charged ions with a minimum signal of 500.0 were selected for high-energy collision-induced dissociation (HCD) and fragmentation via collision-induced dissociation (CID), both using an isolation width of 2.0 m/z. HCD had an activation time of 10 ms and a normalised collision energy of 45%, while CID had the same activation time but a normalised collision energy of 35%.

MaxQuant v2.6.5(Cox and Mann, 2008) was used to search the raw MS data against a protein database obtained by translating genomes (*Lanistes nyassanus*, *P. canaliculata*, *P. maculata*, *P. diffusa*, *M. cornuarietis*, *Pila pesmei*) and transcriptomes (*P. scalaris*) that contained protein sequences (target) and their reversed sequences (decoy). Search parameters were 20 ppm for the first search mass tolerance; 0.5 Da for fragments; two maximum missed cleavages for trypsin. The matched peptides were further filtered to delete reverse and potential contamination using Perseus v2.1.3 (Tyanova et al., 2016). A false discovery rate threshold of < 0.01 was also applied in each replicate for final protein identification. Only proteins that were identified in at least two biological replicates, contained at least 2 unique peptides were retained.

To further confirm the number of PV1 subunit, MSGFPlus searches (https://github.com/MSGFPlus/msgfplus) were performed for the ovorubin MS data of *P. canaliculata* purified by Horacio Heras team, with following parameters: ±20 ppm parent mass tolerance; isotope error range (-ti ‘-1,2’); fully tryptic enzyme settings (-e 1 -ntt 2); conducting a parallel search against a decoy protein database (-tda 1) for calculating the false discovery rate (FDR). Only proteins that were identified in three biological replicates and contained at least 3 unique peptides were retained.

*New genome assemblies reveal the evolution trajectory of PV2 subunits*

Genomic analyses reveal an expanded tachylectin gene family (5–8 copies) in *Pomacea* species—functionally linked to the MACPF chain for embryonic protection (Giglio et al., 2020). Our previous study hypothesized the evolutionary history of two PV2 subunits (Sun et al., 2019), with the updated chromosomal-level genomes of *Pomacea* and *Marisa* (Xiong et al., 2025), we have reconstructed the evolutionary trajectory of PV2. Briefly, we identified 41 MACPF homologs across eight ampullariids and sixteen molluscs. Maximum-likelihood phylogenetics resolved two clades (Fig. S13A): Clade I (12 ampullariid sequences) nested among non-ampullariid sequences, while Clade II (29 ampullariid-specific homologs) underwent extensive duplication in New-World snails. PV2-associated MACPF genes (Pcan_009G001433; Pmac_011G001277) evolved within Clade II, exhibiting albumen gland-specific high expression (TPM > 500) and PVF proteomic detection (Table S15). Divergence dating places their origin at ~6 million years ago, postdating the *P. diffusa* split with *P. canaliculata* and *P. maculata* (Fig. S14). Thus, neofunctionalization after duplication—specifically, MACPF secretion by the albumen gland and partnership with tachylectin—drove PV2 toxin emergence in terrestrial-egg-laying *Pomacea*.

Tachylectin-like genes appear specific to Caenogastropoda, with 66 homologs identified in ampullariids and 11 in *Littorina sinensis* and *Rapana venosa*, but undetected in other molluscan species. Maximum-likelihood phylogenetics divided these into two clades (Fig. S13B): Clade II contains six ampullariid sequences that high expression in albumen gland (one each in *P. scalaris*, *P. maculata*, and *P. canaliculata*, plus three in *M. cornuarietis*; TPM >500), while Clade I comprises 25 ampullariid sequences featuring a terminal *Pomacea*-specific group of 20 genes. Divergence dating revealed that Clade I underwent extensive duplication ~30 Ma after *Pomacea* diverged from other ampullariids (Fig. S16-17), with three *Pomacea* genes (Pcan_009G001432, Pmac_011G001273, Pmac_011G001275) exhibiting both high albumen gland expression and PVF encoding. These originated 6-8 Ma—contemporaneous with PVF-associated MACPF evolution—supporting neofunctionalization after duplication as the mechanism for the novel tachylectin subunit in Canaliculata eggs.

Genome screening reveals that at least one scaffold/chromosome contains both MACPF-like and tachylectin-like genes in all six ampullariids (Fig. S18). This two-gene configuration, present in *Pila celebensis* and *L. nyassanus*, likely existed as a single copy in the ampullariid ancestor. Following the divergence of *M. cornuarietis* and *Pomacea* lineages (~34 Ma; Figs. S14-S17), *M. cornuarietis* acquired species-specific MACPF duplications. Extensive duplications then occurred in *Pomacea* post-divergence, exemplified by at least three tandemly duplicated MACPF-tachylectin pairs on *P. canaliculata* chromosome 9 (Fig. S18), suggesting coordinated duplication events. Within the Canaliculata clade, specific genes (Pcan_009G001433, Pmac_011G001277, Pcan_009G001432, Pmac_011G001273, Pmac_011G001275) underwent positive selection post-*P. diffusa* divergence. These exhibit both high albumen gland expression and MACPF-tachylectin complex binding sites (Giglio et al., 2020), explaining restriction of PV2 within Canaliculata. Overall, the occurrence of PV2 in Canaliculata eggs—and its toxicity—indicates a defensive role against terrestrial predators. The co-selection of PV2 and terrestrial egg deposition represents a key innovation enabling their aquatic-to-terrestrial transition and contributes significantly to the global invasiveness of *Pomacea*.

**2. Supplementary Figs. S1 to S18**

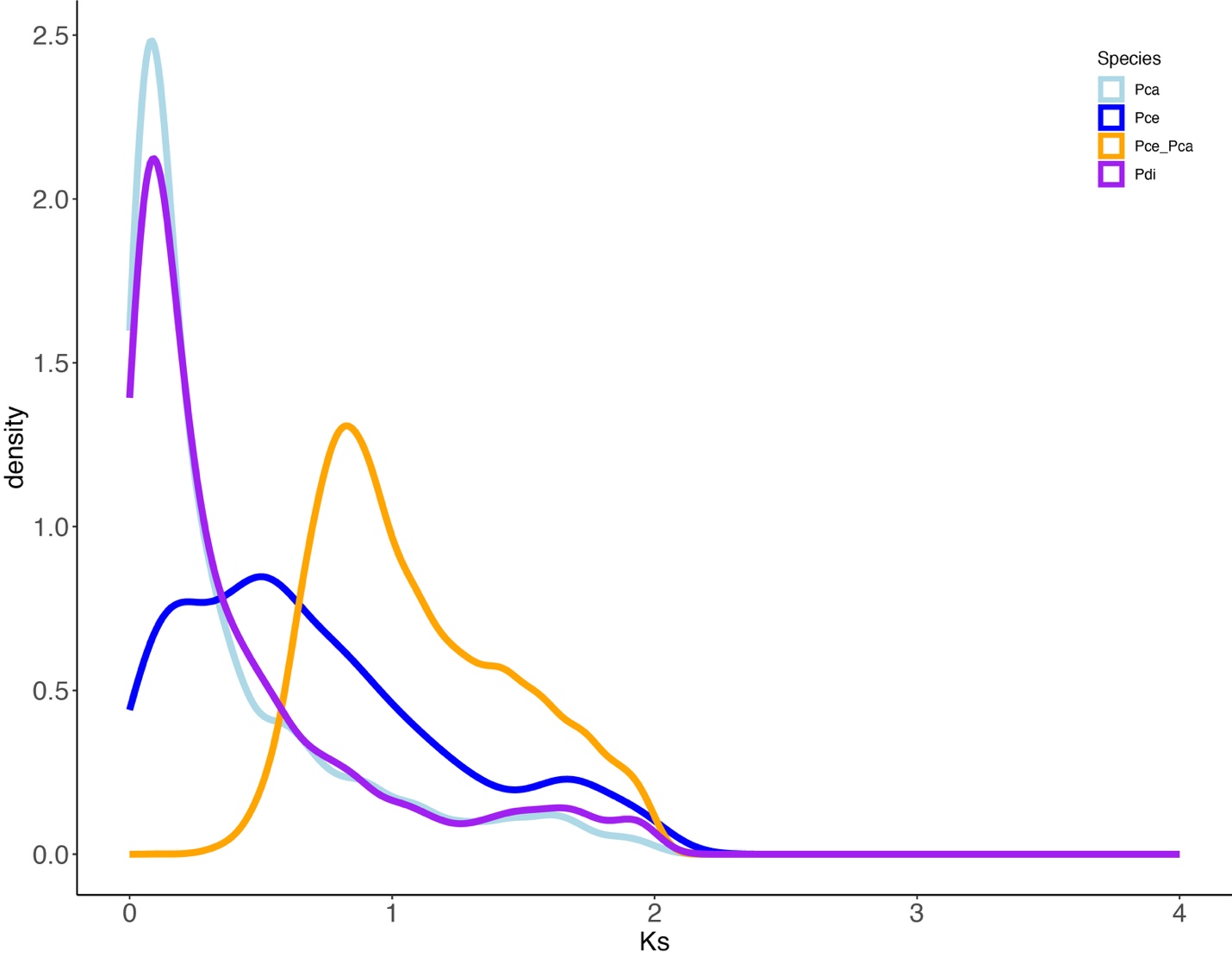

**Fig. S1.** The distribution of the synonymous substitution rate (Ks) of homologous gene groups for intraspecies and interspecies comparisons.

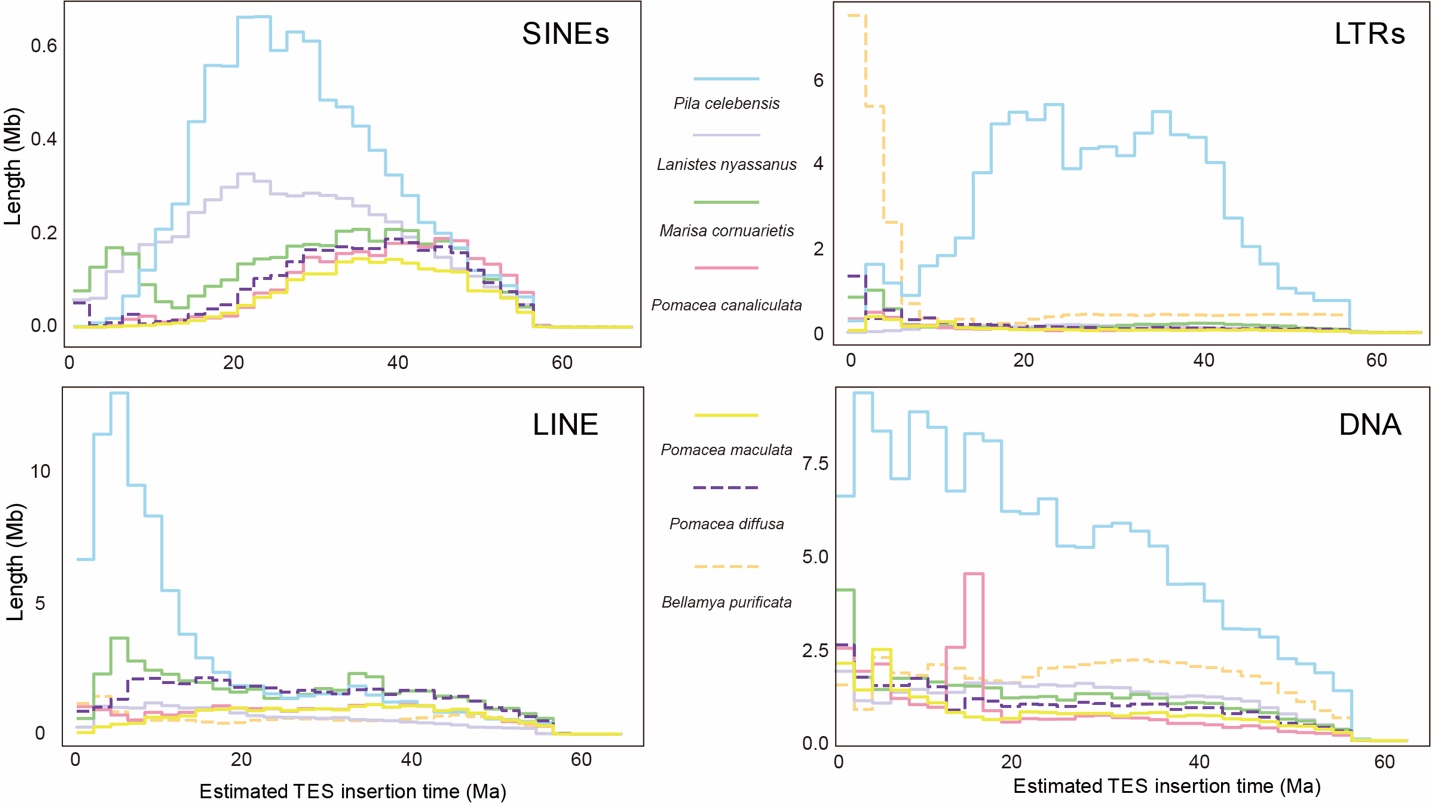

**Fig. S2.** Comparison of estimated insertion times of four transposable element (TE) classes among Ampullariidae and *Bellamya purificata*, showing that the genome of *Pila celebensis* is distinct in that the major bursts of SINEs and LTRs corresponded to the time of the split between *Pila* and *Lanistes* (20-40 Ma).

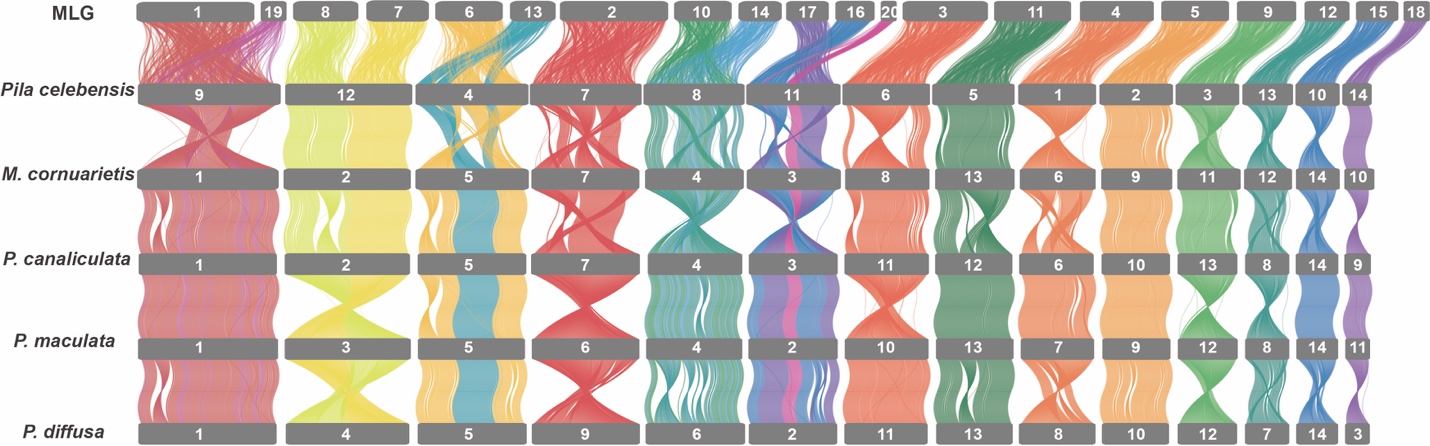

**Fig. S3.** Syntenic rearrangements of molluscan linkage groups (MLGs) within the evolution of ampullariid genomes, with conserved syntenic regions, genome rearrangements, and structural variations among these species in relation to the ancestral genome organization.

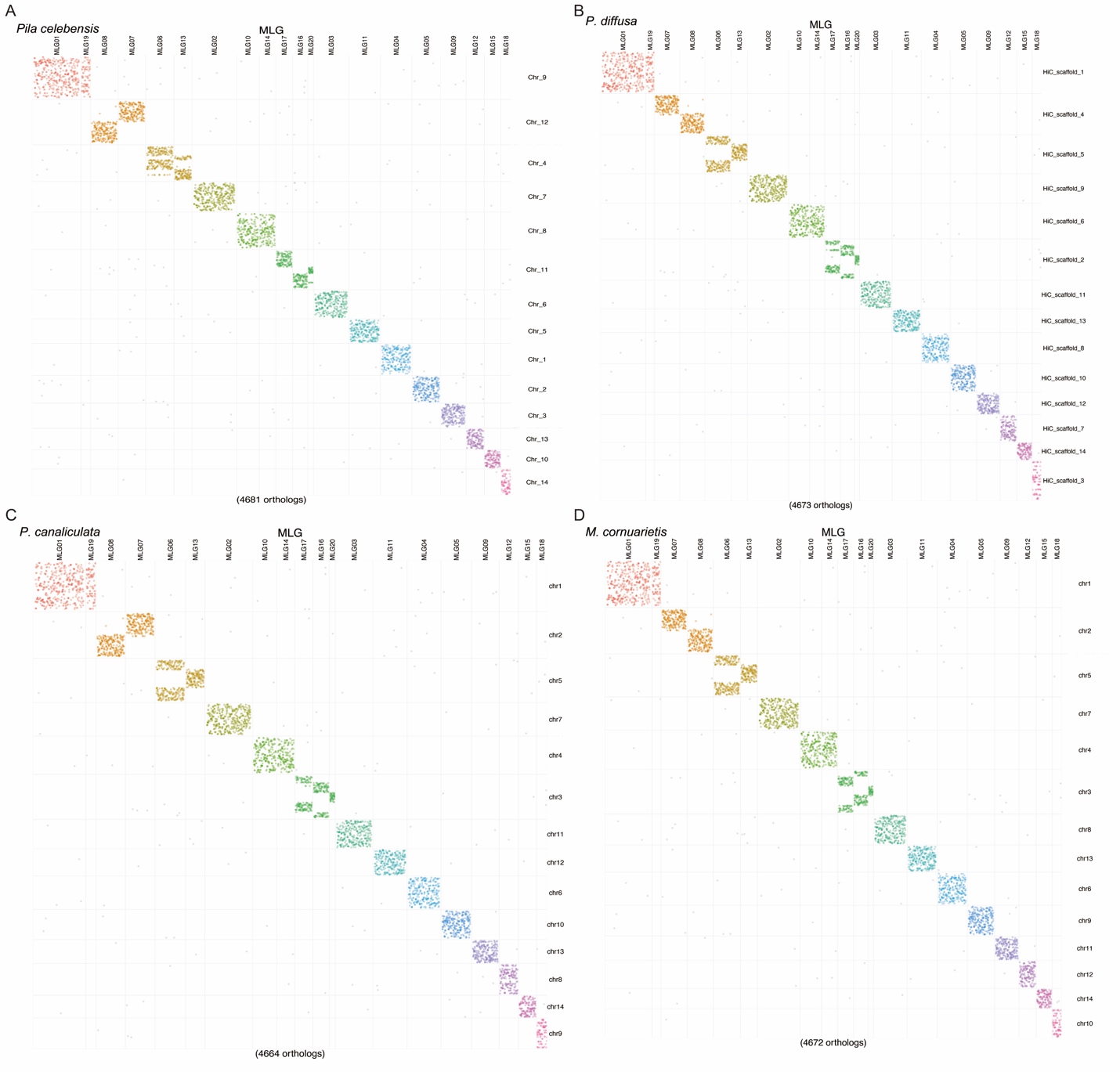

**Fig. S4.** Oxford dotplots of the chromosome-scale ancient gene linkage with four Ampullariidae species, *Pila celebensis*, *Pomacea diffusa*, *Pomacea canaliculata*, *Marisa cornuarietis*.

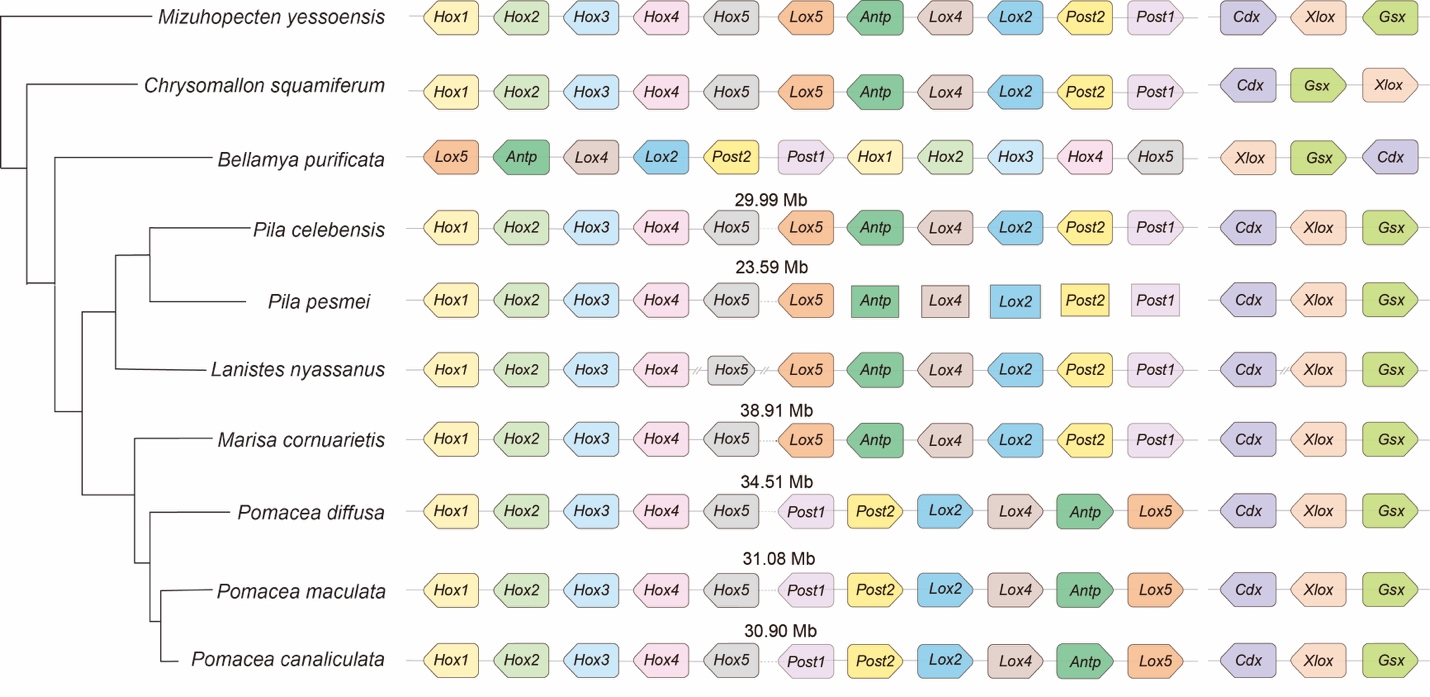

**Fig. S5.** Schematic illustration of *Hox* and *ParaHox* gene clusters in seven ampullariid species and three other selected molluscans. Gene names are labeled in center of the colored graphs. Genes on the same scaffold or chromosome are connected with a line, but the line length does not represent sequence length. The symbol “//” indicates a break between different scaffolds. Transcription direction, when available, is indicated by a bullet head.

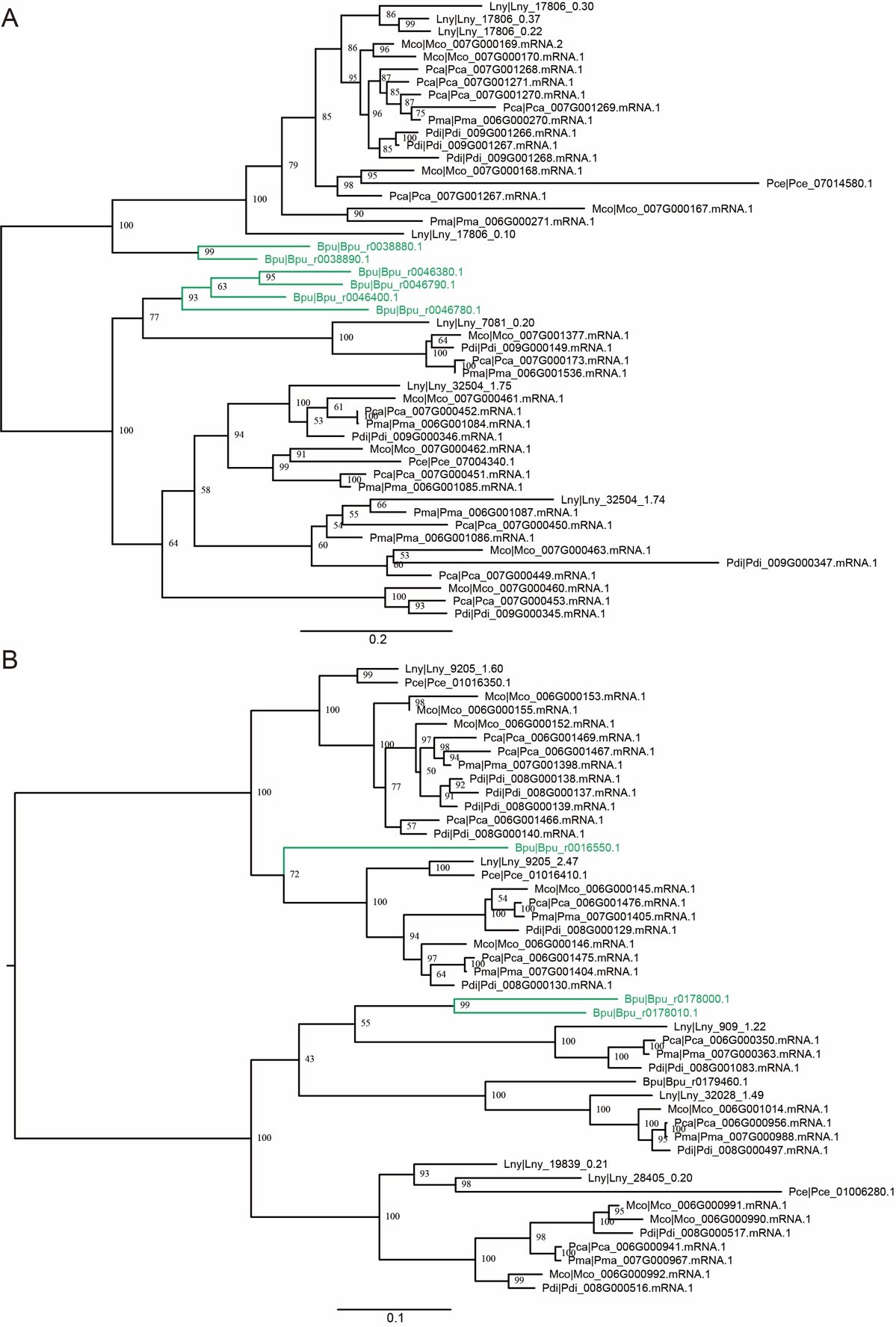

**Fig. S6.** Phylogencitc trees of expanded gene families in Ampullariidae. **(A)** Cellulase gene family and **(B)** β-D-xylosidase gene family. The green branches indicate the outgroup of Ampullariidae – *B. purificata*.

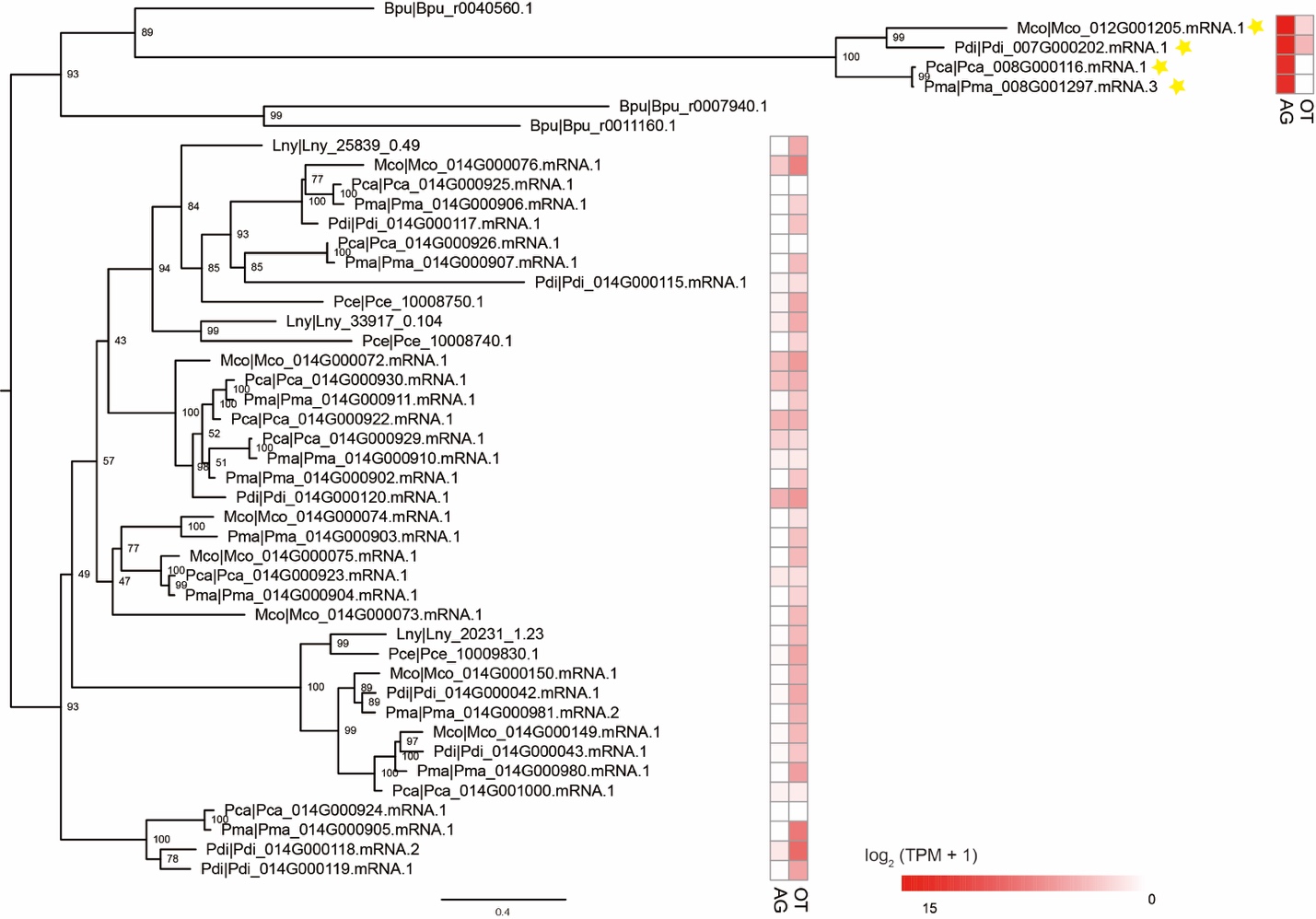

**Fig. S7.** Phylogencitc trees of expanded C1q-related gene family in Ampullariidae. Expression of these genes in the albumen gland (AG) and other tissues (OT) are showed on the right hand side, yellow star indicate PVF proteins identified in PVF MS data (Table. S13).

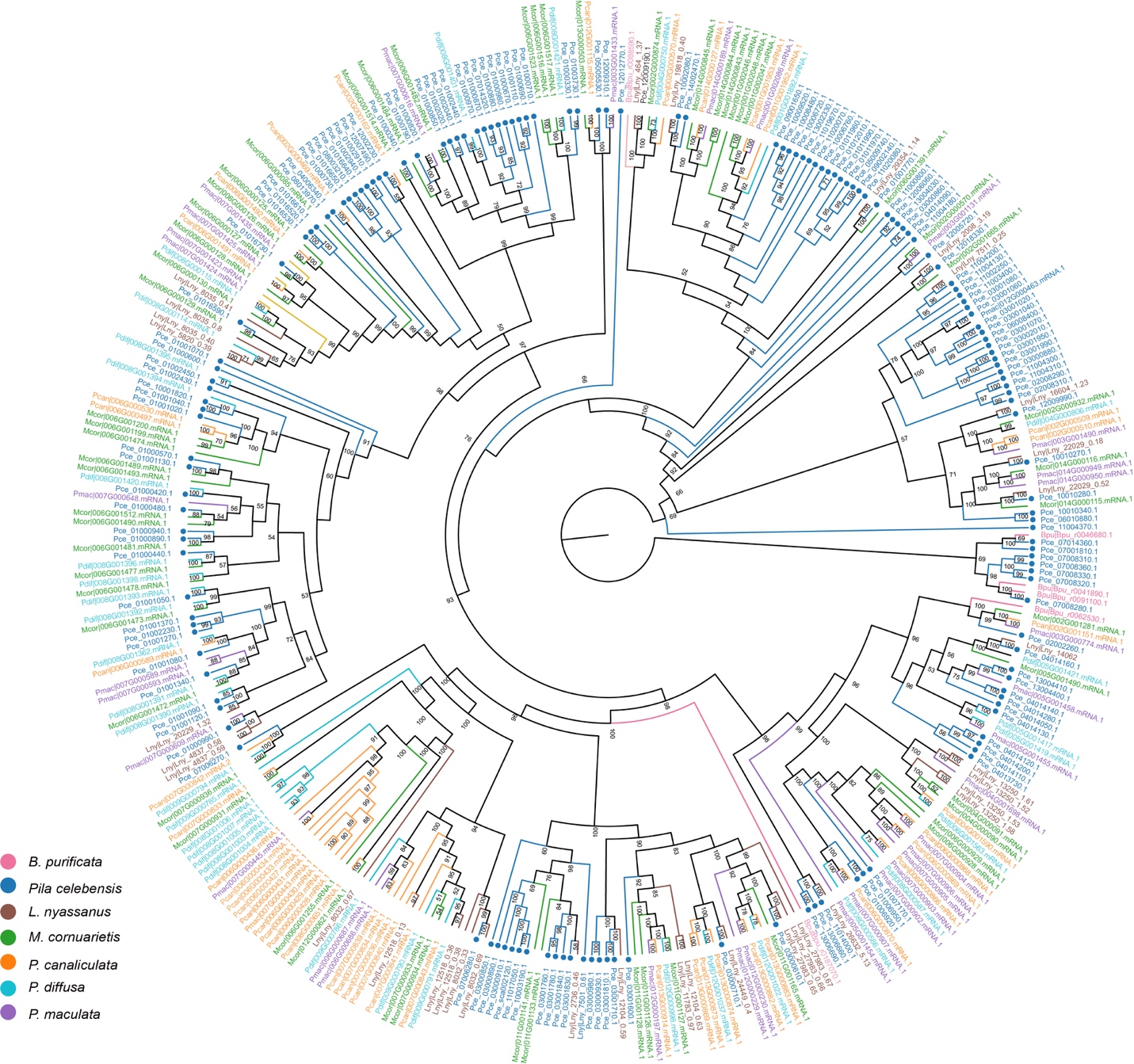

**Fig. S8.** Unrooted maximum-likelihood tree showing the massive expansion of the G-protein coupled receptor (GPCR) family in Ampullariidae and *Bellamya purificata*.

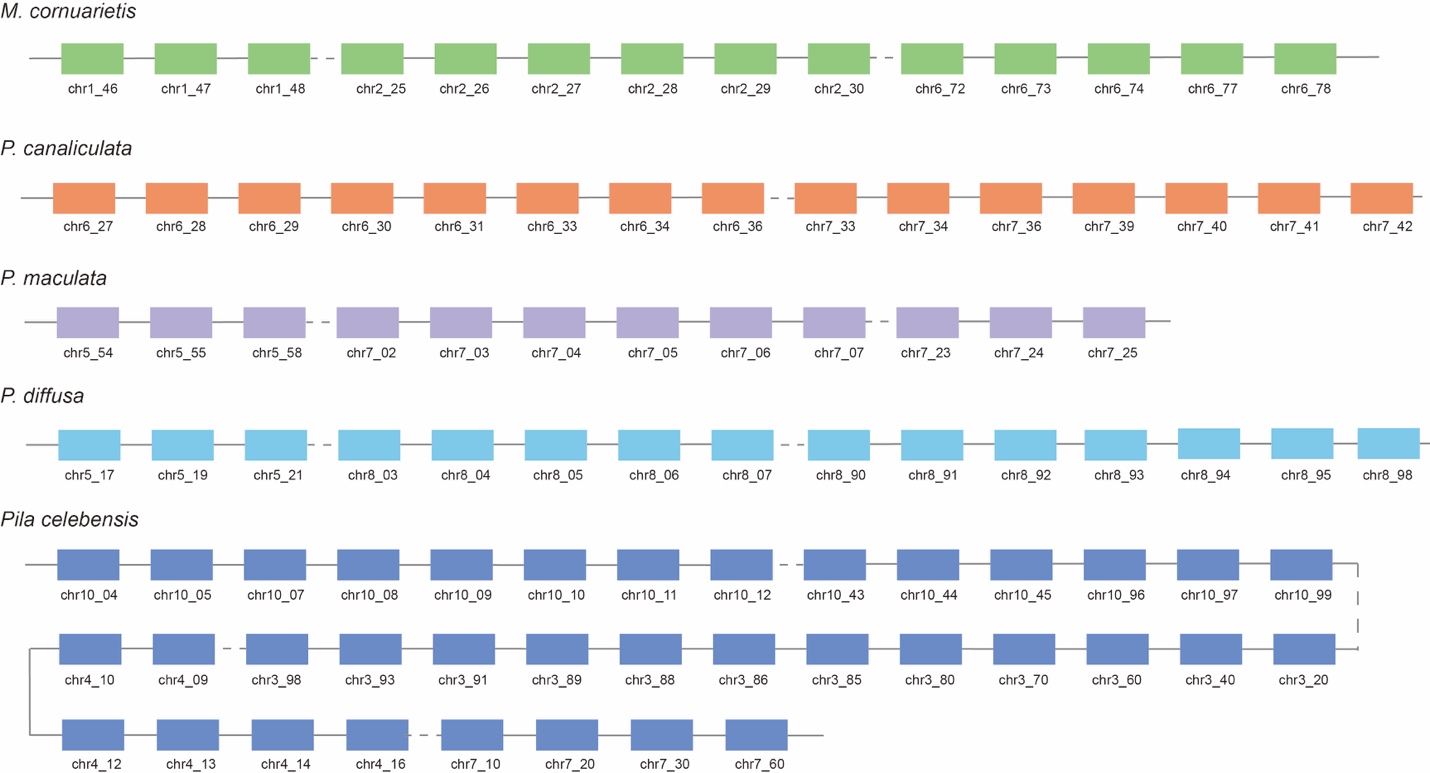

**Fig. S9.** Schematic illustration of GPCR genes in Ampullariidae. Gene names are labeled under the colored graphs and the line length does not represent sequence length.

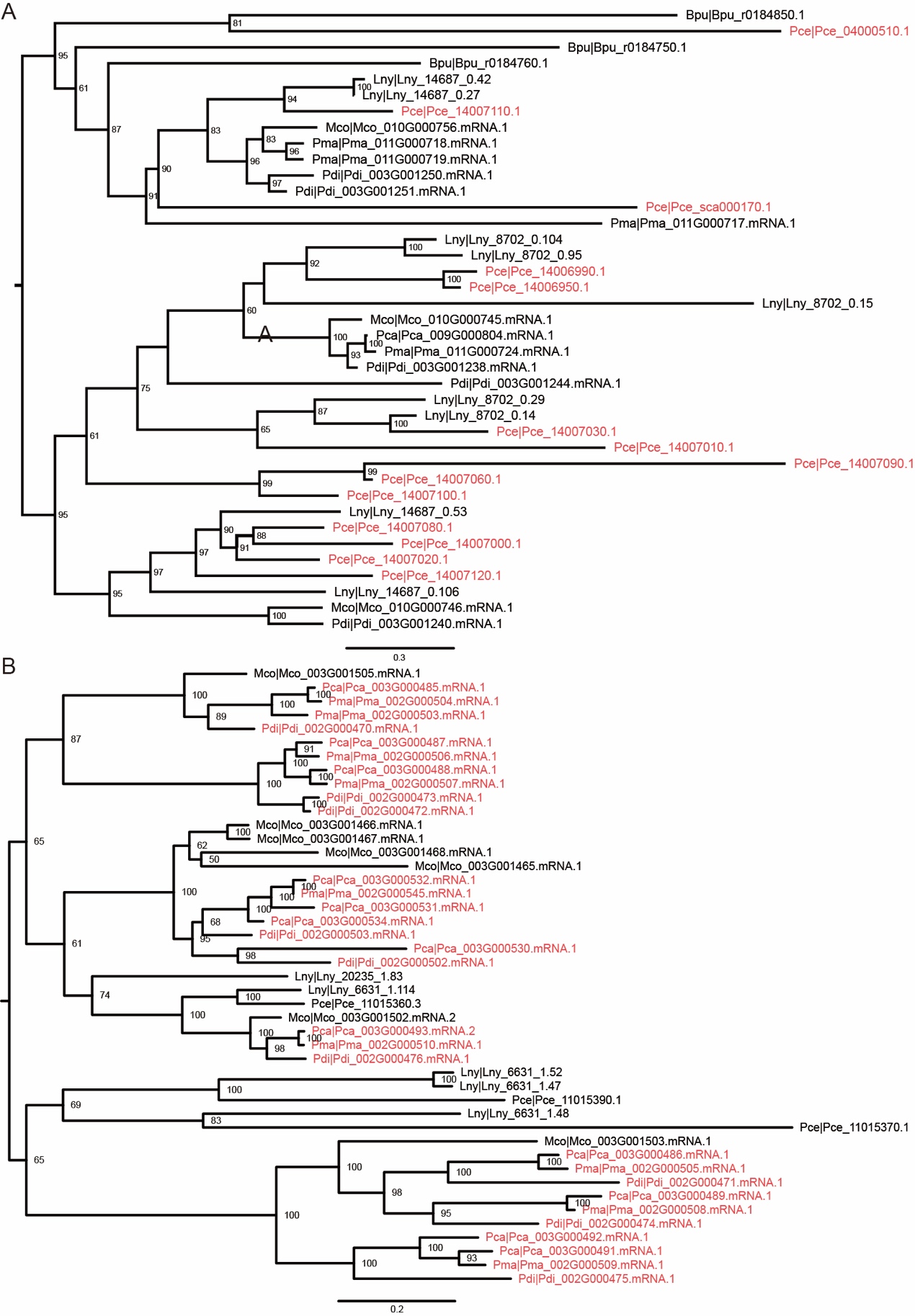

**Fig. S10.** Phylogencitc trees of lineage-specific expanded gene families in *Pila* and *Pomacea* respectively. **(A)** keratin gene family and **(B)** neprilysin gene family. The red branches indicate the expanded genes in *Pila* and *Pomacea*.

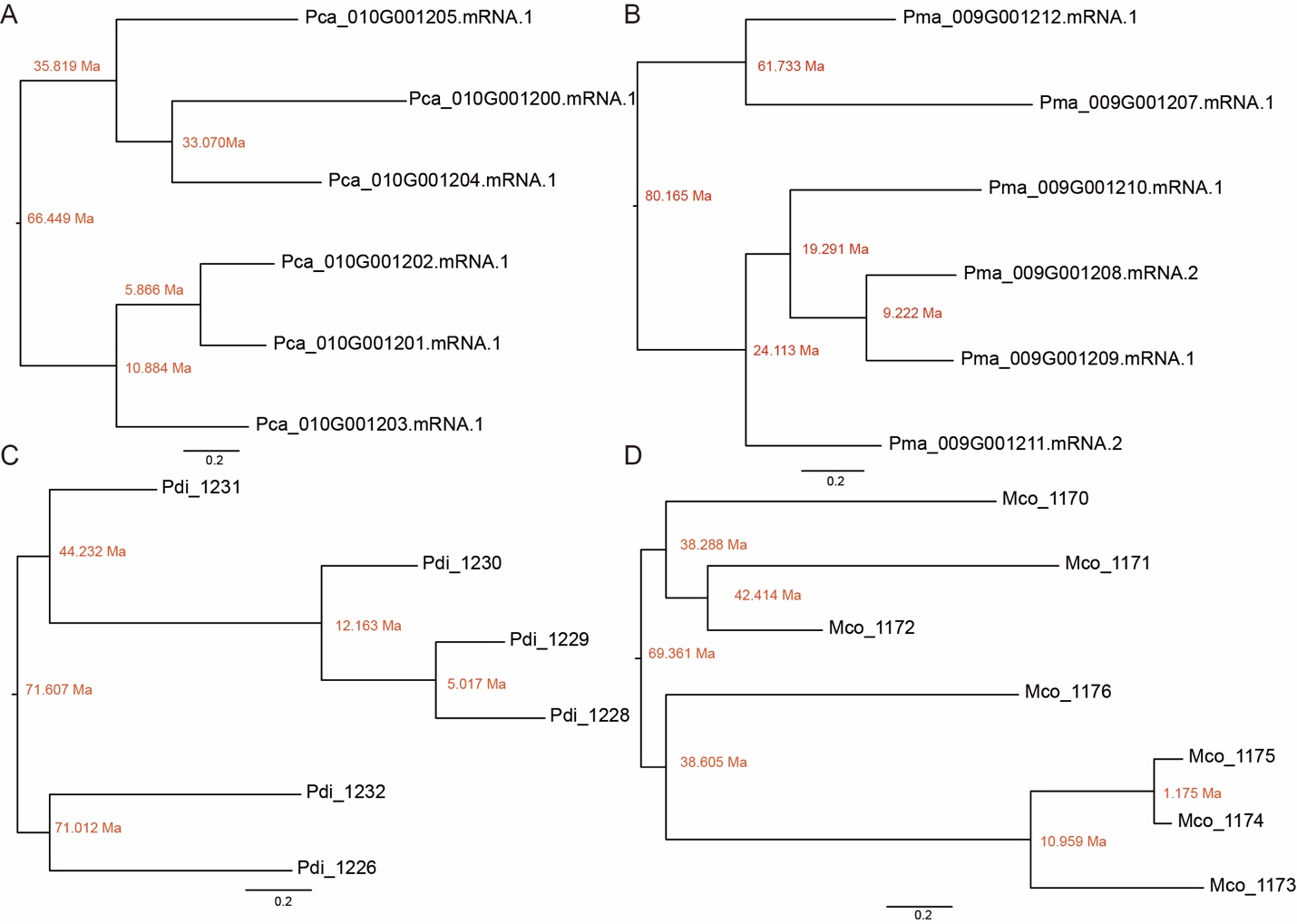

**Fig. S11.** Divergence time of PV1 sequences in four New world species in Ampullariidae. **(A)** *Pomacea canaliculata*, **(B)** *P. maculata*, **(C)** *P. diffusa* and **(D)** *Marisa cornuarietis*. Sequences within the red rectangle belong to clade V of PV1, and the other sequences belong to other four clades. The red number next to nodes are divergent time calculated by KaKs_calculator.

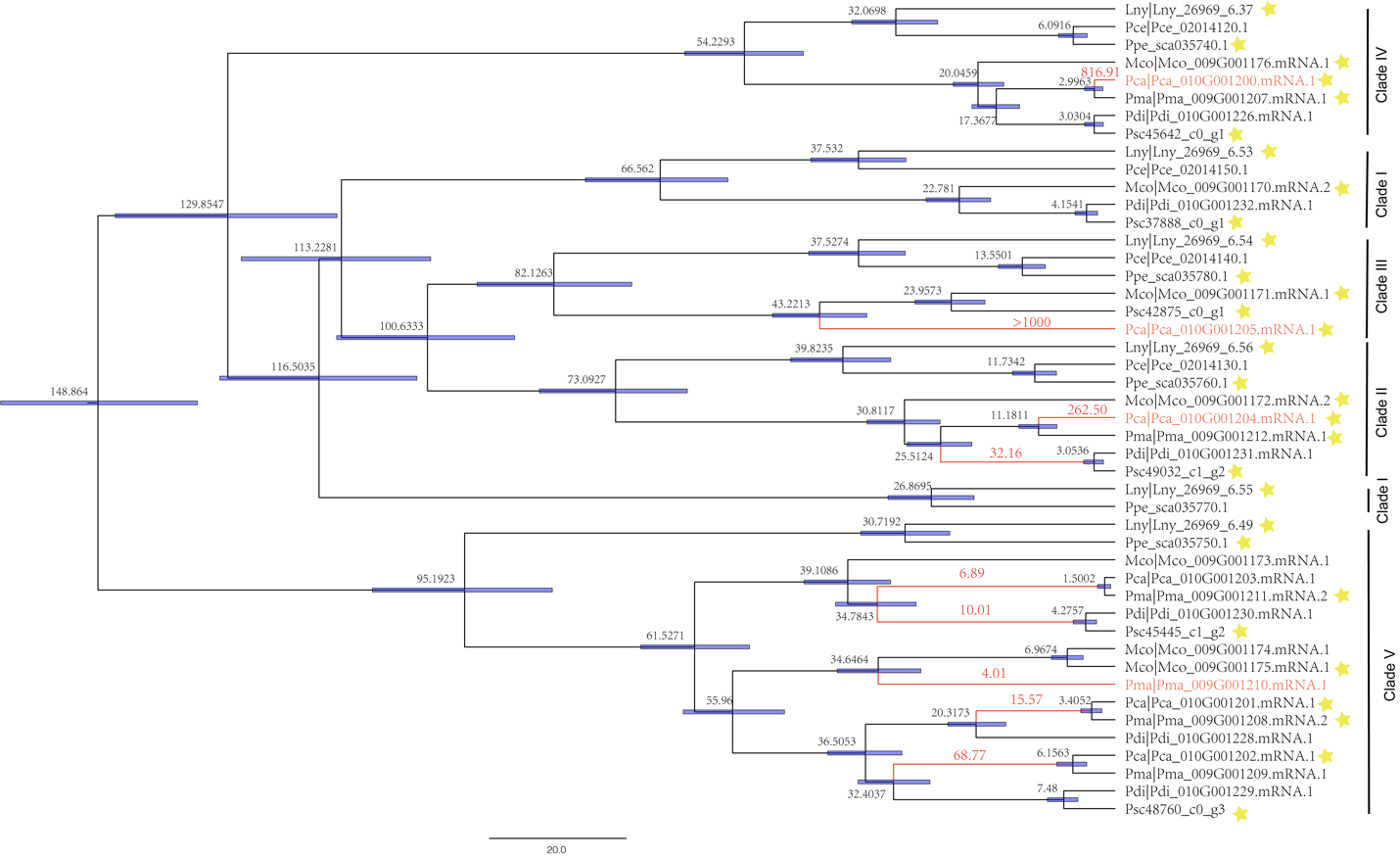

**Fig. S12.** Maximum-likelihood phylogenetic relationships among PV1 sequences in ampullariidae. The tree was calibrated at root node using fossils events to reveal divergence times. Blue lines indicate 95% confidence interval for divergence times. Red branch means under positive selection and the red number on branches are dN/dS values. The yellow star indicate the protein detected in PVF MS data.

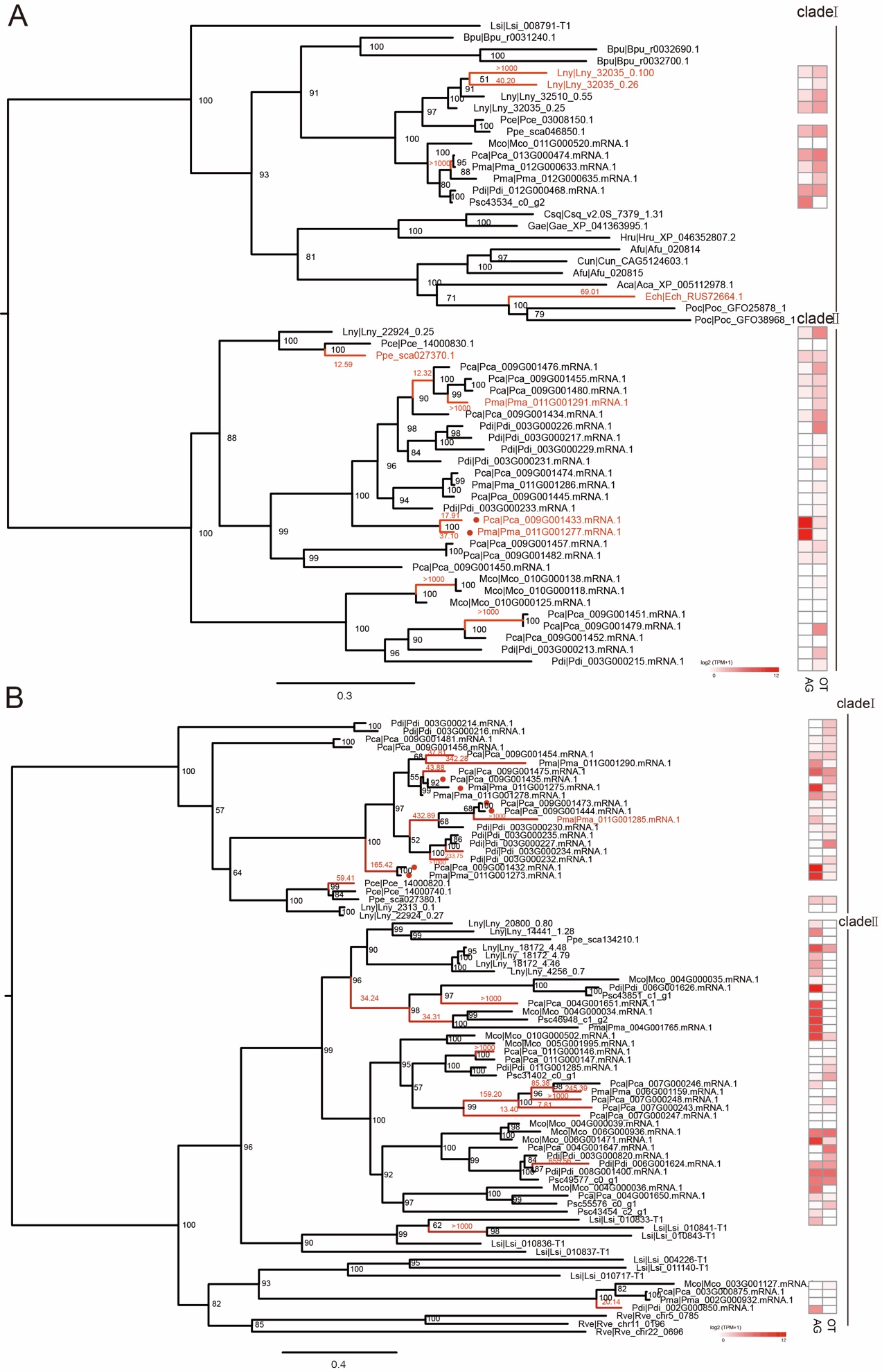

**Fig. S13.** Phylogeny and evolution of PV2 genes in ampullariidae. Phylogeny and expression of homologues of (A) MACPF-like genes and (B) tachylectin-like genes. Numbers on nodes are bootstrap values (>50%) and the red dot preceding each sequence name indicates detection of these proteins in the mass spectrometry (MS) dataset. Red branch means under positive selection and the red number on branches are dN/dS values. Gene expression levels are in logarithmic scale. AG, albumen gland; OT, other tissues.

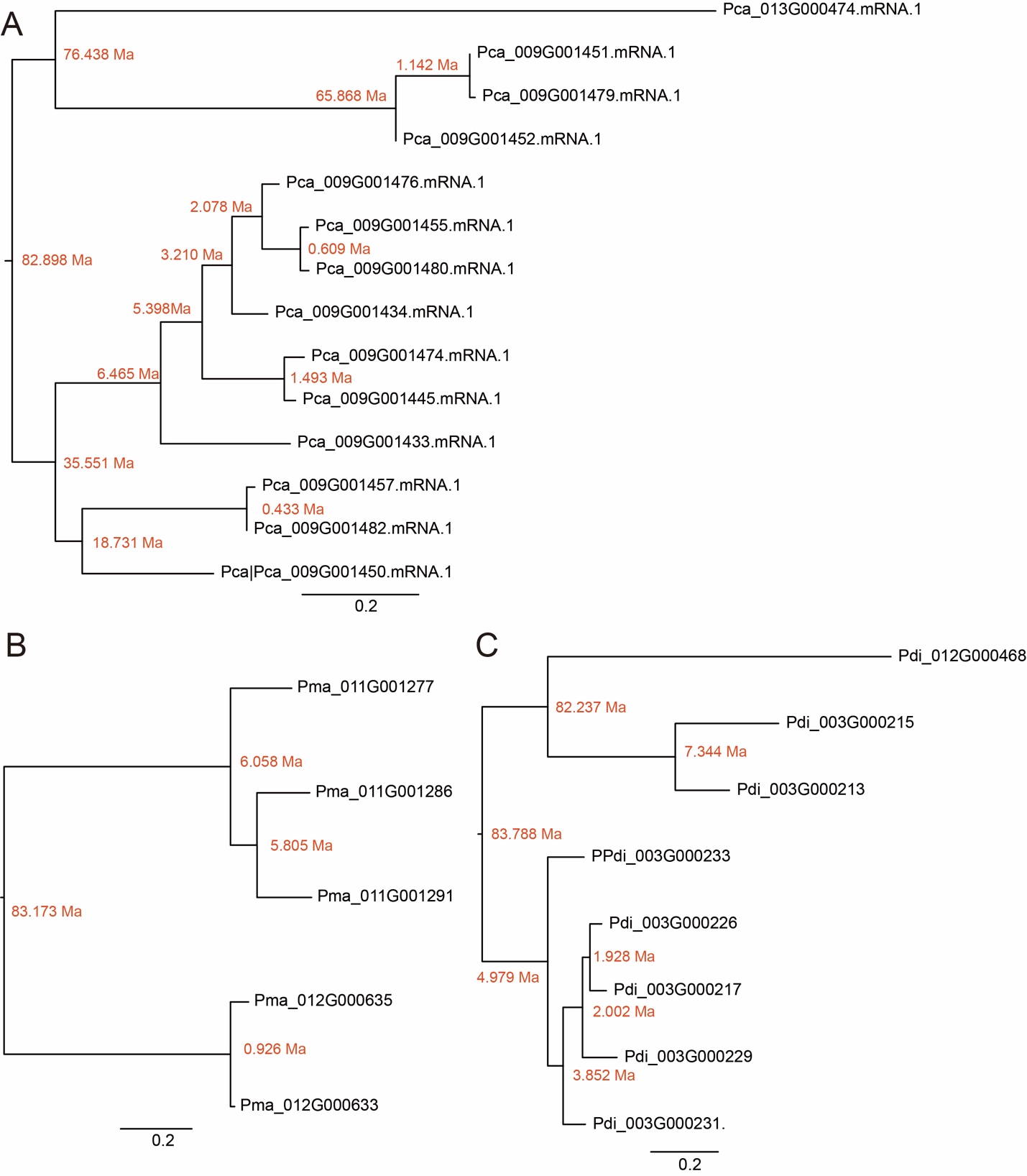

**Fig. S14.** Divergence time of MACPF-like sequences in three New-world species in Ampullariidae. **(A)** *Pomacea canaliculata*, **(B)** *P. maculata*, and **(C)** *P. diffusa*. Sequences within the red rectangle belong to Clade I of MACPF, and other sequences belong to Clade II of MACPF. The red number next to nodes are divergent time calculated by KaKs_calculator.

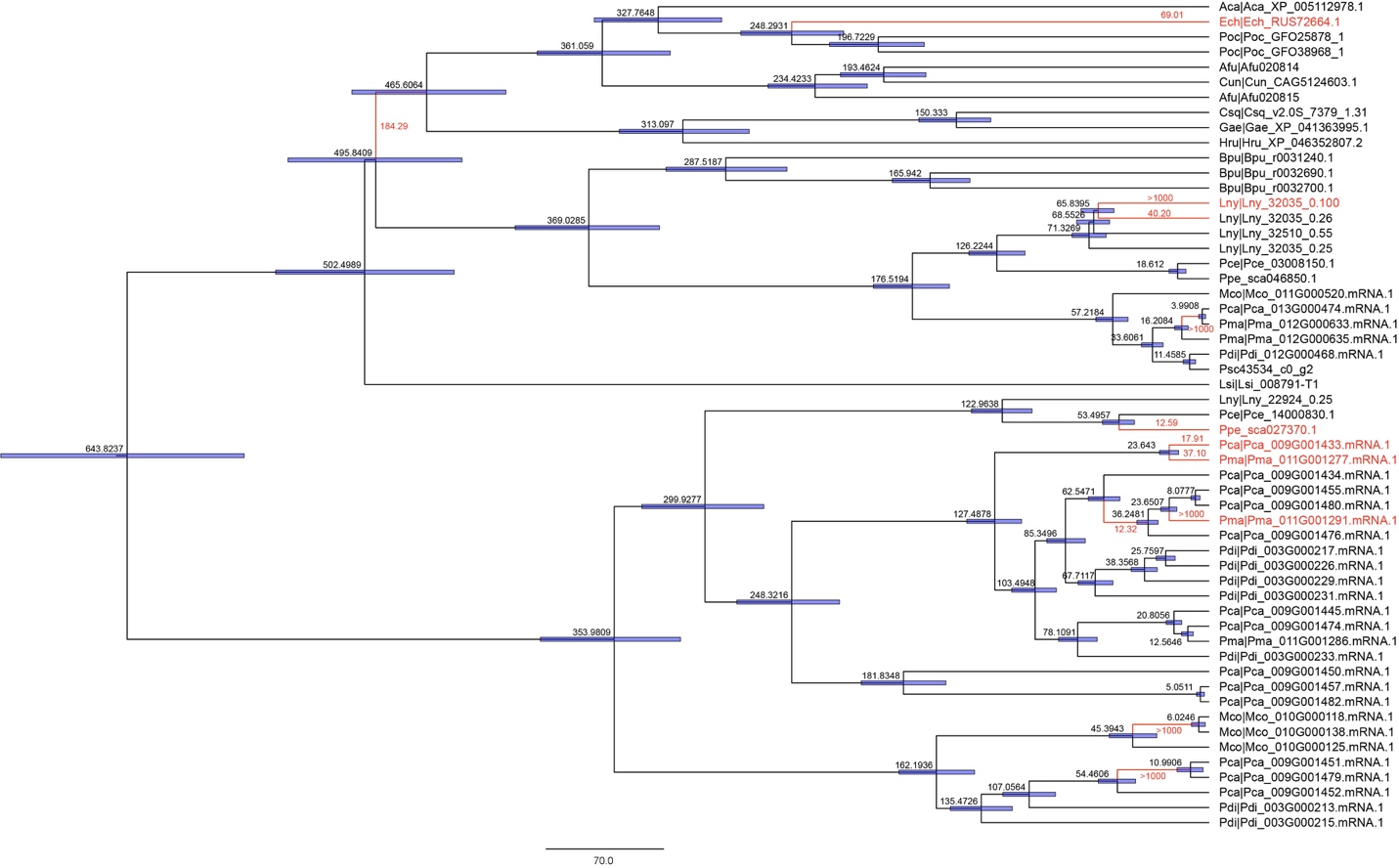

**Fig. S15.** Maximum-likelihood phylogenetic relationships among MACPF sequences in 23 molluscs. The tree was calibrated using fossils events to reveal divergence times. Blue lines indicate 95% confidence interval for divergence times. Red branch means under positive selection and the red number on branches are dN/dS values.

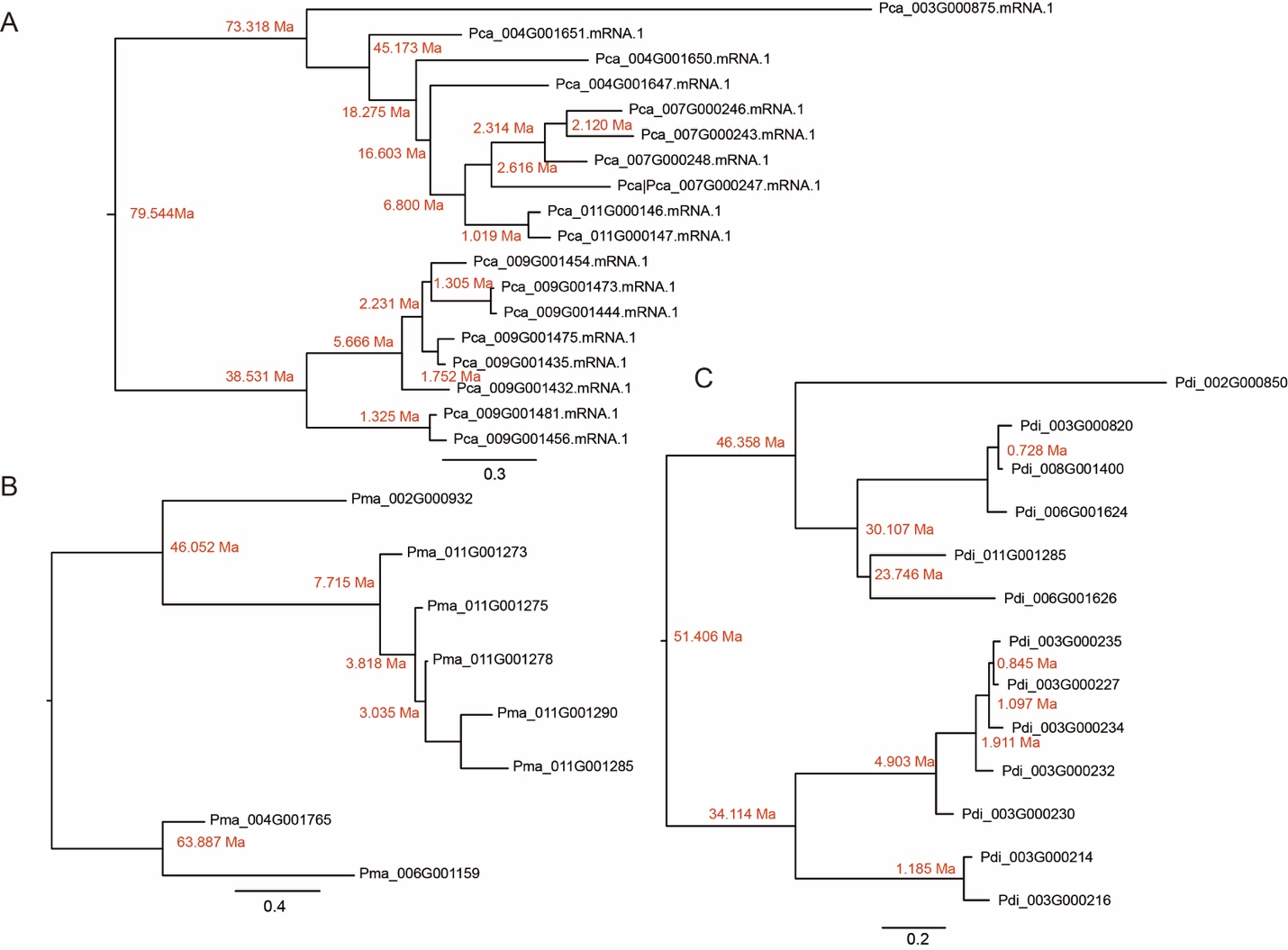

**Fig. S16**. Divergence time of tachylectin-like sequences in three New-world species in Ampullariidae. **(A)** *Pomacea canaliculata*, **(B)** *P. maculata*, and **(C)** *P. diffusa*. The red number next to nodes are divergent time calculated by KaKs_calculator.

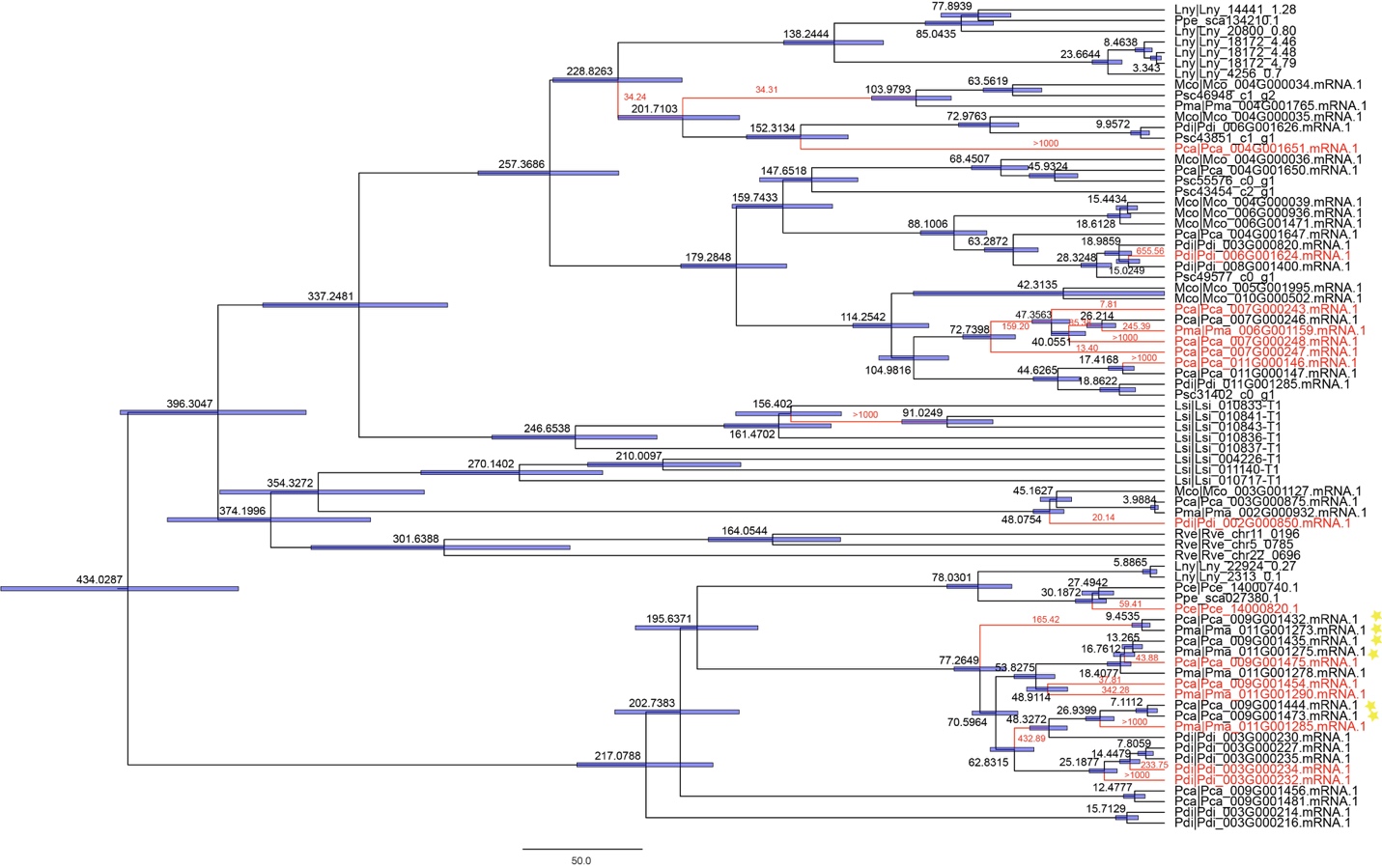

**Fig. S17.** Maximum-likelihood phylogenetic relationships among tachylectin sequences in Caenogastropoda. The tree was calibrated using fossils events to reveal divergence times. Blue lines indicate 95% confidence interval for divergence times. Red branch means under positive selection and the red number on branches are dN/dS values. Yellow star indicates the sequence was detected in protein MS dataset.

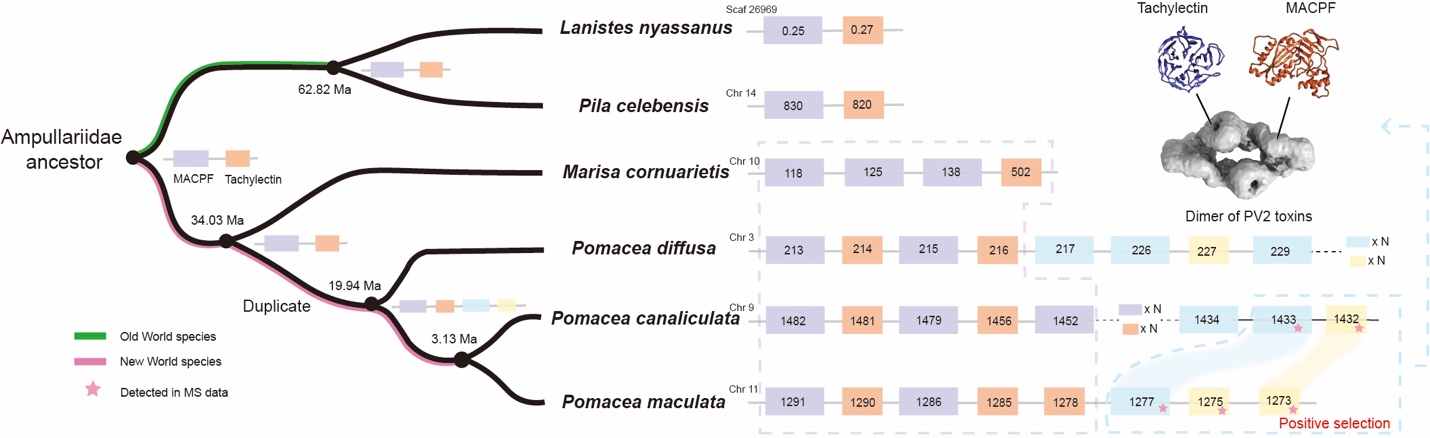

**Fig. S18.** Schematic of PV2 evolution. A two-gene configuration (purple and orange section) indicates the common ancestor of PV2 in Ampullariidae. The MACPF and tachylectin genes framed by a blue dotted line underwent positive selection and contain protein-binding sites for MACPF-tachylectin complexes (Giglio et al., 2020). Pink star indicates highly expressed in the albumen gland.
